# Conformation-specific Antibody Deciphers K27-linked Ubiquitination in Chaperone-Mediated Proteostasis

**DOI:** 10.64898/2025.12.18.695067

**Authors:** Chengxiao Han, Yicheng Weng, Qingyun Zheng, Qian Qu, Satchal K. Erramilli, Zhen Su, Yujuan Duan, Yunxi Han, Xiaoguo Zhai, Jingxian Li, Anthony A. Kossiakoff, Man Pan, Minglei Zhao, Lei Liu, Yuanyuan Yu

## Abstract

Lysine 27 (K27)-linked polyubiquitination plays critical yet incompletely defined roles in proteostasis, innate immunity, and disease progression; however, investigations into this process have long been hindered by its extremely low abundance and the lack of conformation-specific enrichment tools. Herein, we describe the development of a long-sought conformation-specific antibody, K27-IgG, which can selectively recognize—among all ubiquitin chain types—the unique architecture of K27-linked polyubiquitin (K27-polyUb) characterized by a distinct buried K27-isopeptide bond, with high affinity (KD = 4.66 nM). This antibody was derived from synthetic antibodies initially generated via phage display, using chemically synthesized K27-linked diubiquitin (K27-diUb) as the antigen. High-resolution co-crystal structures uncovered the unique K27-diUb interface targeted by these sAbs. Subsequent reformatting of these sAbs into a full-length human immunoglobulin G (IgG) scaffold yielded K27-IgG, notably exhibiting markedly enhanced affinity without compromising selectivity. Using K27-IgG as a tool, we achieved sensitive detection and immunoprecipitation (IP) of endogenous K27-polyUb in cells, and delineated the intracellular interaction landscape of K27-polyUb through complementary proteomic approaches. Two key findings emerged: 1) The molecular chaperone DNAJB1 is a specific reader of K27-linked ubiquitin chains (but not other linkages) and that K27-polyUb chains themselves exhibit chaperone-like activity, suggesting a novel mechanism by which K27-polyUb regulates chaperone-mediated proteostasis; 2) The E2 enzyme UBE2Q1 assembles K27-diUb, identifying it as a potential writer for this ubiquitin chain topology. Collectively, this study establishes K27-IgG as a robust tool for deciphering the K27-linked ubiquitin code, thereby opening new avenues for investigating the biological functions of K27-linked polyubiquitination.

**Highlights:**

- First K27-linkage conformation-specific antibody with nanomolar affinity overcomes a major barrier in the field.
- K27-IgG unlocks functional mapping of the K27 ubiquitin landscape under proteotoxic stress.
- Molecular chaperone DNAJB1 is a selective “reader” of K27-linked ubiquitin chains.
- K27 chains possess intrinsic chaperone activity, enabling protein refolding and suppressing aggregation.
- E2 enzyme UBE2Q1 is a “writer” that directly assembles K27-linked ubiquitin chains.

**Graphical abstract:** 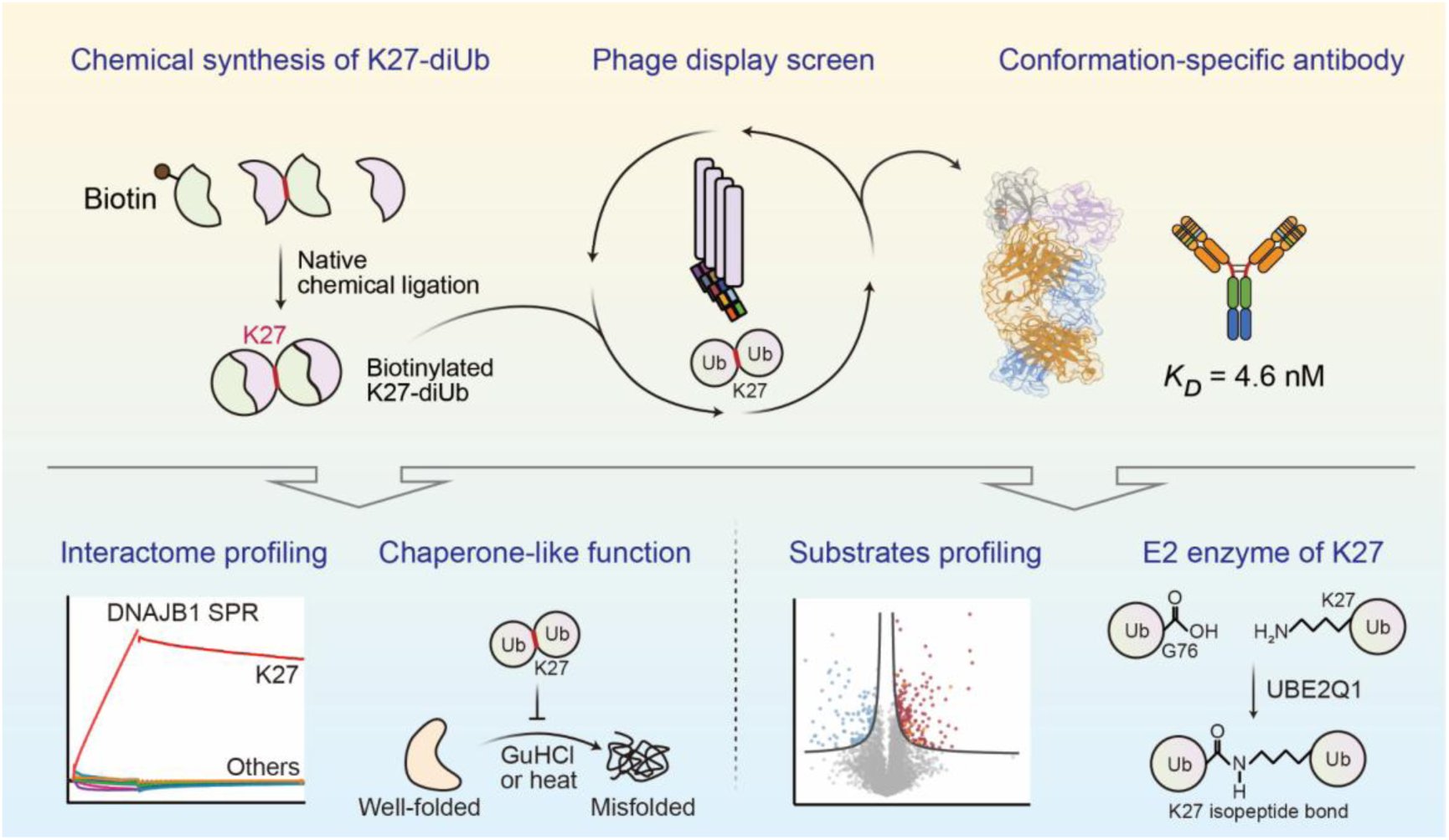

## Introduction

Ubiquitination is one of the most versatile and dynamic post-translational modifications, acting as a master regulator of diverse cellular processes including protein degradation, DNA repair, immune signaling, and cell cycle progression^1–4^. This remarkable functional diversity arises from ubiquitin (Ub)’s unique structural plasticity. Its C-terminal carboxyl group can form isopeptide bonds with any of eight distinct amino groups (M1, K6, K11, K27, K29, K33, K48, or K63) on another Ub, thereby generating polyUb chains with distinct architectures. Each linkage adopts a unique three-dimensional topology (Figure 1A) that dictates specific interactions with Ub-binding domains, forming sophisticated “ubiquitin codes” that precisely tailor the recruitment of effector proteins to trigger appropriate cellular response^1,3^.

**Figure 1.**
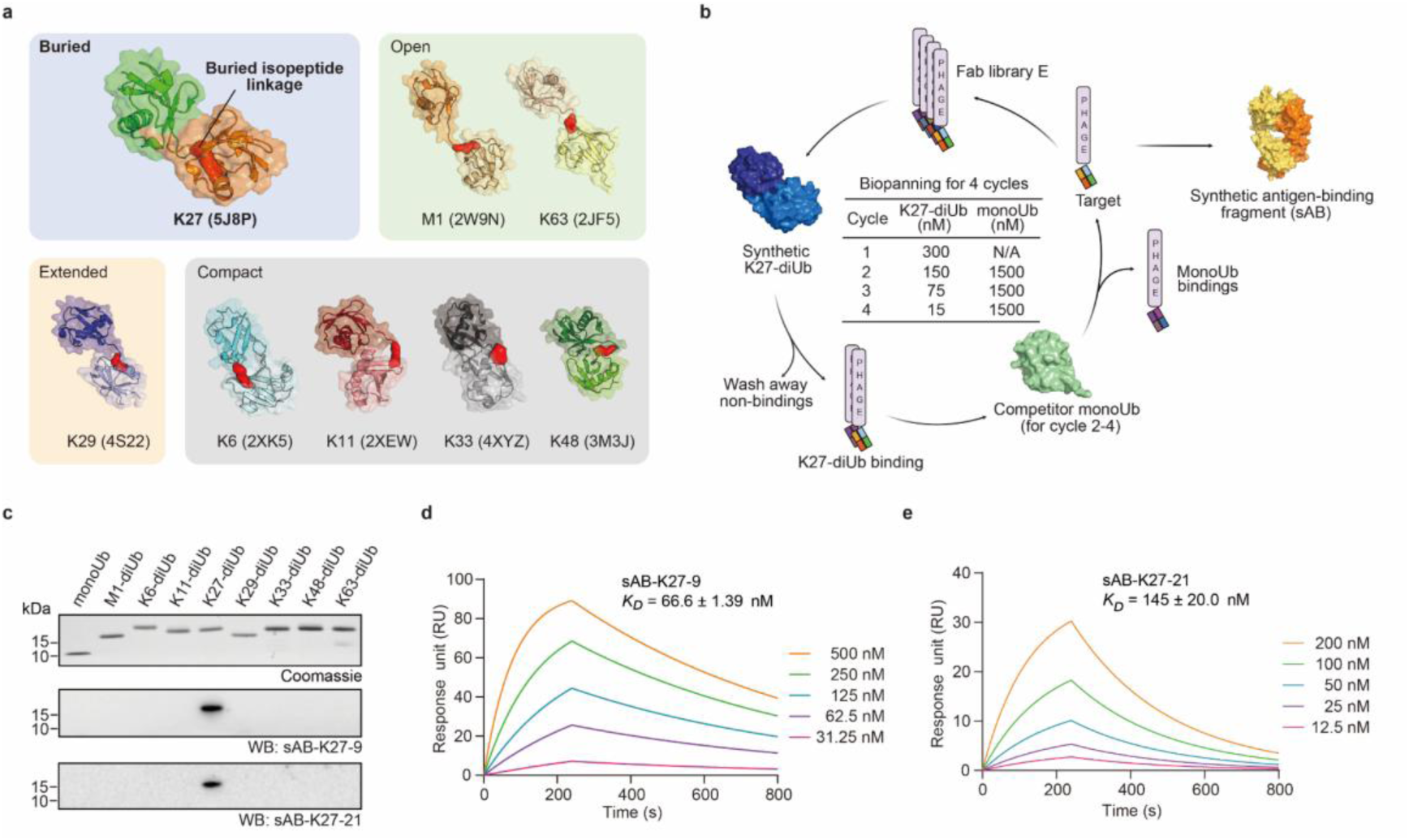
Selection and characterization of conformation-specific synthetic antibody fragments for K27-linked diUb. **(a)** Crystal structures of diUb with eight linkages: K27 (PDB 5J8P), M1(PDB 2W9N), K6 (PDB 2XK5), K11 (PDB 2XEW), K29 (PDB 4S22), K33 (PDB 4XYZ), K48 (PDB 3M3J), and K63 (PDB 2JF5). Isopeptide bond surfaces are highlighted in red. **(b)** Schematic overview of phage display selection strategy used to isolate conformation-specific sABs targeting biotinylated K27-linked diUb. **(c)** Western blot analysis of sAB-K27 specificity against eight synthetic diUb linkages. 100 ng of synthetic diUb were loaded into each lane. **(d-e)** Surface plasmon resonance (SPR) analysis of sAB-K27-9 (d) and sAB-K27-21 (f) binding to K27-linked diUb. Dissociation constants (*K_D_*) are shown as mean ± standard deviation (s.d.) (n = 3). All gel panels in this figure are representative at least three independent experiments.

Dysregulation of this precise ubiquitin code is increasingly implicated in human pathologies. K48-linked chains typically direct substrates for proteasomal degradation, and defects in their assembly contribute to neurodegenerative disorders such as Parkinson’s and Alzheimer’s diseases^5–9^. K63-linked ubiquitination orchestrates DNA repair pathways and immune responses, with impairments driving cancer development^10–13^. Emerging evidence suggests that atypical linkages (e.g., K27 and K29) are key regulators of immune disorders and metabolic diseases^14–26^. Notably, despite their low abundance^27–30^, K27-linked chains have been implicated in diverse processes, including proteostasis^31^, autophagy^32–36^, innate immunity^20,21,37–39^, DNA damage repair^40^, tumorgenesis^22,33,34,41,42^, and Parkinson’s disease^16,32,43^. This functional versatility positions K27-linked chains as a crucial yet understudied component of cellular signaling networks.

The linkage-specific organization of Ub chains serves as both a core regulatory mechanism and a disease-relevant node, making decipherment of the ubiquitin code a central priority in ubiquitin research^1,2,44–46^. Linkage-specific antibodies with conformation-selectivity have advanced understanding of distinct Ub linkages. For example, K48- and K63-specific antibodies clarified their roles in RIP1- and IRAK1-dependent innate immune signaling^47^; a K11-specific antibody identified the E3 ligase APC/C as a “writer” mediating mitotic degradation^48^; an M1-specific antibody revealed M1-linked chains’ key role in NF-κB signaling^49^; a K6-specific affimer revealed HUWE1 as the primary K6 “writer”^50^; a K11/K48 bispecific antibody revealed branched chains’ dual functions in mitosis regulation and rapid proteasomal clearance^51^; a K29-specific synthetic antigen-binding fragment (sAB) highlighted K29-linked chains’ roles in proteotoxic stress responses and cell cycle regulation^15^; and nanobodies targeting K48/K63 branched chains identified VCP/p97-related proteins as binders and debranching enzymes^52^. Despite these advances, K27-linked polyubiquitination features a structurally unique Ub chain with its isopeptide bond buried in a solvent-inaccessible interface^29,30,53^—presenting a unique challenge for developing K27 conformation-selective antibodies to dissect its cellular functions.

Using phage display technology with chemically synthesized K27-diUb as the antigen, we isolated two conformation-selective synthetic antibodies (sAB-K27-9 and sAB-K27-21) that exhibit exclusive selectivity for K27-linked ubiquitin chains. High-resolution co-crystal structures of sAB-K27-K27-diUb complexes uncovered the molecular basis of linkage-specific recognition. Notably, reformatting sAB-K27-21 into a full-length human IgG (K27-IgG) markedly enhanced performance, achieving ∼30-fold higher binding affinity (*K_D_* = 4.66 nM). K27-IgG enabled nanogram-level detection of K27-polyUb in biochemical assays and immunostaining-based imaging. Leveraging K27-IgG’s specificity, we used complementary native and denaturing immunoprecipitation-mass spectrometry (IP-MS) to map the intracellular K27-linked Ub landscape. Our integrated approach revealed two key findings: (i) the molecular chaperone DNAJB1 is a specific K27-polyUb interactor, indicating K27-polyUb’s regulatory role in chaperone-mediated proteostasis, and (ii) the E2 enzyme UBE2Q1 assembles K27-diUb, identifying it as a potential “writer” for this Ub chain topology. These results establish K27-IgG as a powerful and versatile tool that opens new avenues for investigating the K27 ubiquitin code.

## Results

### Selection and characterization of K27-linkage specific sAB

Our work began with the chemical synthesis of K27-linked diUb. To facilitate subsequent synthetic antibody (sAB) selection and characterization, we incorporated either an N-terminal biotin or His_6_-tag into the distal ubiquitin. In addition, a double 2-(2-(2-aminoethoxy)ethoxy)acetic acid (AEEA) linker was inserted between biotin and the diUb moiety to enhance flexibility. We successfully prepared K27-linked diUb using a route that combines solid-phase peptide synthesis (SPPS) and hydrazide-based native chemical ligation (NCL) (Figure S1A)^30,54,55^. The integrity of the chemically synthesized K27-linked diUbs was verified via high-performance liquid chromatography (HPLC) and liquid chromatography-mass spectrometry (LC-MS) analyses (Figure S1B-C). Following gradient guanidine-HCl (GuHCl) dialysis, the folded K27-diUb was purified by size-exclusion chromatography (SEC) and validated using sodium dodecyl sulfate-polyacrylamide gel electrophoresis (SDS-PAGE) (Figure S1D). Circular dichroism (CD) spectroscopy showed structural fingerprints matching those of monoubiquitin (monoUb), confirming the native folding of synthesized K27-diUb (Figure S1E).

We next performed phage display panning of Library E, a humanized Fab-based synthetic antibody library^56^, against K27-linked diUb, using monoUb as a competitive binder to enhance selection specificity (Figure 1B). After several rounds of iterative panning with progressively increased stringency^57^, we successfully isolated two high-affinity sAB clones, designated sAB-K27-9 and sAB-K27-21.With these two sABs in hand (Figure S2A), we systematically evaluated their specificity across all eight ubiquitin linkage types (Figure 1C). Both sABs showed no detectable cross-reactivity with other linkages even under prolonged exposure conditions, confirming their exclusive recognition of K27-diUb. Given the close proximity of the K27 and K29 sites (a potential source of cross-reactivity), we performed western blotting using serial dilutions of K27-linked diUb (50-3.13 ng), alongside 250 ng of K29-linked diUb as a control. Both sABs detected K27-linked diUb at levels as low as 3.13 ng without cross-reactivity to K29-linked diUb (Figure S2B). Binding affinities were quantified by surface plasmon resonance (SPR), giving *K_D_* values of 66.6 ± 1.39 nM for sAB-K27-9 (Figure 1D) and 145 ± 25.0 nM for sAB-K27-21 (Figure 1E).

### Structural characterization of sAB-K27 recognizing K27-linked diUb

To elucidate the molecular basis for conformation-specific recognition, we determined the crystal structures of sAB-K27 in complex with K27-linked diUb. These complexes were SEC-purified (Figure S3A) and verified by SDS-PAGE analysis (Figure S3B). The stable complex was subjected to crystallization screening (see Methods for details). The crystal structures were resolved at 2.2 Å for sAB-K27-9 and 2.9 Å for sAB-K27-21 (Table S1).

Both structures revealed a 1:1 stoichiometry between sAB-K27 and K27-linked diUb. For the interaction between sAB-K27-9 and diUb (Figure 2A), the hairpin-like CDR-H3 (residues E106, Y107, W108, Y110, Y111, Y112, and W115) was sandwiched between distal Ub (residues K6, I44, H68 and R72) and proximal Ub (residues I44 and H68). The CDR-H1 (residues Y33, S34 and Y5) make contact with distal Ub (residues R42, Q49 and R72), while the CDR-L3 (residues W93 and W94) interacted with the proximal Ub (residues L8, V70 and L71) (Figure 2B). On the Ub side, the I44 patches of both Ub units were engaged in modular interactions with the hairpin-like CDR-H3 (Figure S3C). For the interaction between sAB-K27-21 and K27-linked diUb, a similar binding mode was observed (Figure S3D). Notably, CDR-H3 adopted a more rigid β-hairpin conformation at the central interface (Figure S3E), which may enhance the structural rigidity of sAB-K27-21.

**Figure 2.**
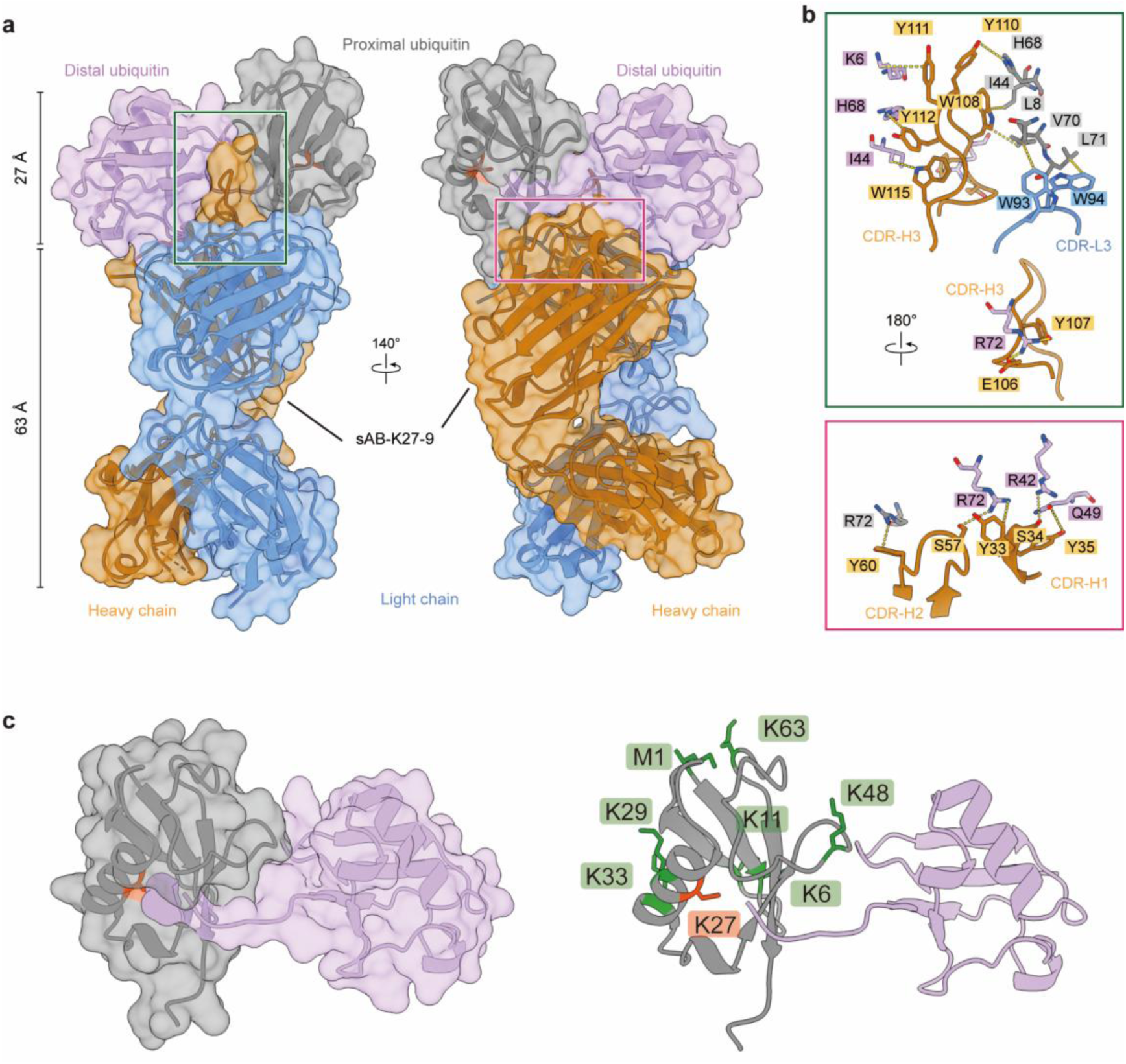
Crystal structures of sAB-K27 in complex with K27-linked diUb. **(a)** Crystal structure of the sAB-K27-9-K27-linked diUb complex resolved at 2.2 Å. The heavy chain (orange) and light chain (blue) of sAB-K27-9 engage the distal (pink) and proximal (grey) ubiquitin units. The isopeptide bond is highlighted in red. **(b)** Detailed views of the binding interfaces between sAB-K27-9 and K27-linked diUb. Putative hydrogen bonds are shown as dashed lines. **(c)** A close-up view of the proximal Ub reveals that, among the eight lysine residues, only Lys27 is spatially accessible to the C-terminal tail of the distal ubiquitin, thereby enabling K27-specific chain assembly. The K27 isopeptide bond is highlighted in red, while the other lysine side chains in proximal Ub are shown in green.

In both structures, the conformation of K27-linked diUb was highly consistent, with a main-chain root mean square deviation (RMSD) of 2.7 Å, indicating that this topology is unique for K27-linked Ub chains (Figure 2C). A closer view revealed that among the eight amino acid residues on the acceptor ubiquitin, only the residue of lysine 27 can be accessed by the C-terminus of the distal ubiquitin unit (Figure 2D), suggesting a unique conformation adopted by K27-linked diUb.

Furthermore, structural alignment of the proximal ubiquitin unit in our K27-linked diUb structure with both apo K27-diUb (PDB 5J8P) (Figure S3F) and the UCHL3-K27-diUb complex (PDB 6ISU) (Figure S3G) revealed that the isopeptide bond remains solvent-inaccessible—even with substantial rotation of the donor Ub. This finding further highlights the structural uniqueness of K27-linked ubiquitination and underscores the need to develop conformation-selective antibodies.

### Characterization of K27-IgG for detection and enrichment

To address the limited binding affinity of our sABs for K27-linked diUb, we engineered full-length human immunoglobulin variants K27-IgG-9 and K27-IgG-21 to optimize their functional properties for downstream biochemical assays (Figure S4A). With the IgG variants isolated, we measured their binding affinities. Notably, K27-IgG-21 exhibited a ∼30-fold improvement in affinity (*K_D_* = 4.66 ± 0.08 nM) (Figure 3A), whereas K27-IgG-9 showed only a modest change (*K_D_* = 53.8 ± 9.59 nM) (Figure 3B). We next validated their specificity by western blotting against all eight diUb linkages (Figure 3C). Both K27-IgGs selectively recognized K27-linked diUb without detectable cross-reactivity to any other linkage, even under long exposure conditions.

**Figure 3.**
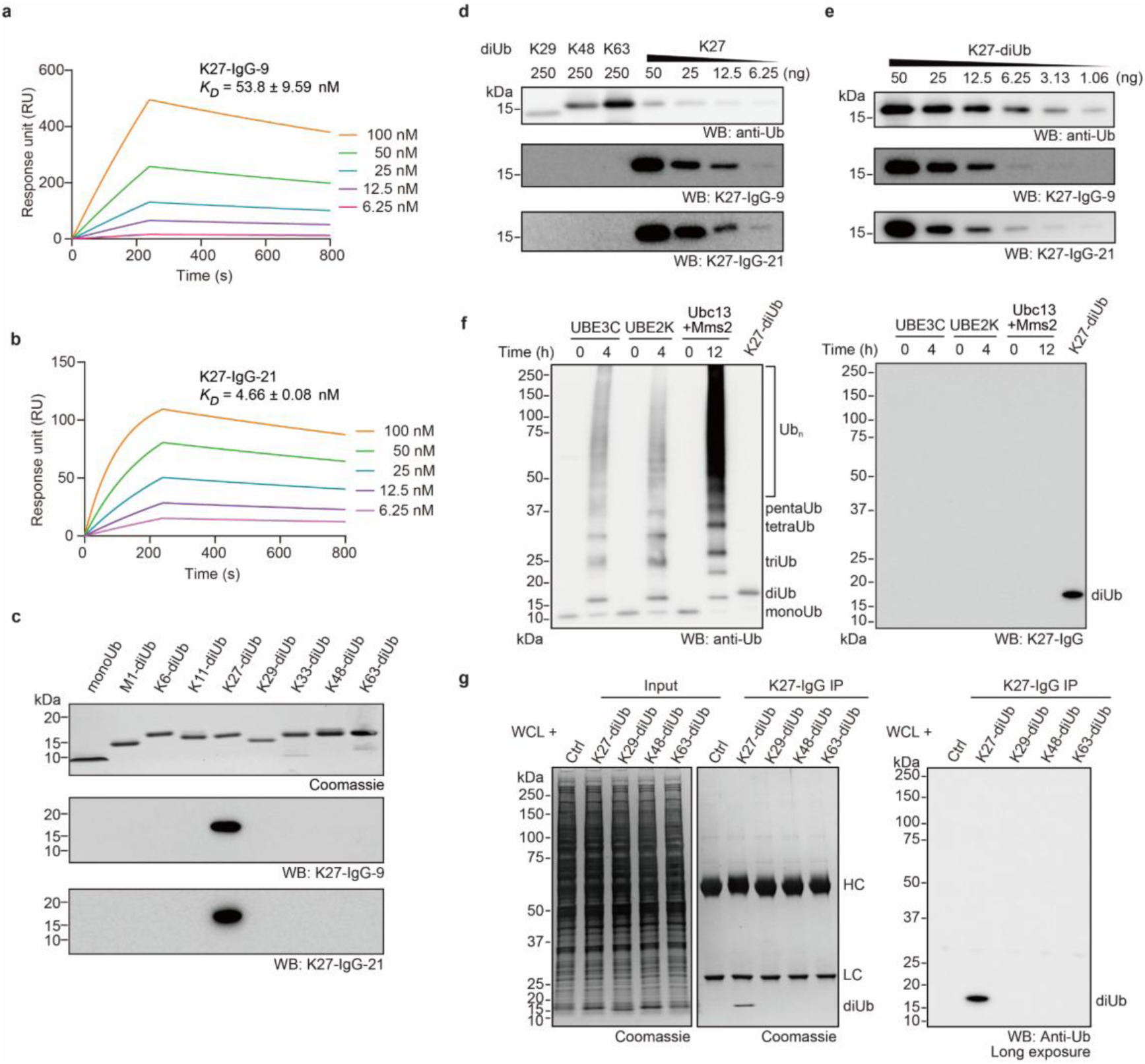
Characterization of K27-IgG for detection and enrichment. **(a-b)** SPR analysis of K27-IgG-9 (a) and K27-IgG-21 (b) binding to K27-linked diUb. *K_D_* are shown as mean ± s.d. (n = 3). **(c)**Western blot of K27-IgG specificity across all eight diUb linkages. 100 ng of diUb was loaded into each lane. **(d)** Western blot demonstrating the specificity of K27-IgG for K27-linked diUb over K29-, K48-, and K63-linked diUbs. **(e)** Western blot assessing the sensitivity of K27-IgG to K27-linked diUb. **(f)** Western blot demonstrating the specificity of K27-IgG for K27-linked diUb over enzymatically assembled K29-, K48-, and K63-linked polyubiquitin. Approximately 1 μg of total ubiquitin was loaded in each lane. **(g)** Immunoprecipitation of K27-linked diUb from HEK293T cell lysates using K27-IgG-21, followed by western blot with an anti-ubiquitin antibody. All gel panels in this figure represent at least two independent experiments.

To rigorously evaluate potential cross-reactivity with more abundant K29, K48, and K63 linkages, we performed western blotting with serial dilutions of K27-linked diUb (50-6.25 ng) alongside high loads (250 ng) of K29-, K48-, and K63-linked diUbs. Remarkably, both antibodies detected K27-diUb at levels as low as 6.25 ng, with no detectable cross-reactivity (Figure 3D). We further assessed the detection limit for both IgGs: K27-IgG-21 could detect K27-linked diUb down to 1.06 ng, whereas K27-IgG-9 exhibited slightly lower sensitivity (detection limit = 3.13 ng) (Figure 3E).

Given its enhanced affinity and sensitivity, K27-IgG-21 was further tested against the K29-, K48-, and K63-linked polyUb chains, which were prepared using UBE3C (mainly K29 and K48^18^), UBE2K (K48^58^), and Ubc13/Mms2 (K63^59^), respectively. Consistent with prior results, K27-IgG-21 selectively recognized K27-linked diUb, with no cross-reactivity to other polyUb chains even under prolonged exposure (Figure 3F; Figure S4B-C).

We further examined applicability of K27-IgG-21 in immunoprecipitation (IP) assays using HEK293T whole-cell lysates supplemented with K27-, K29-, K48-, and K63-linked diUb. As expected, IP results showed robust specific enrichment of K27-linked diUb by K27-IgG-21, with no detectable enrichment of other diUb linkages (Figure 3G). For comparison, we also assessed a commercial sequence-specific antibody targeting K27-linked Ub chains. In direct western blotting, this commercial antibody showed cross-reactivity with K48-diUb (Figure S4D) —consistent with prior reports of its limited specificity^17^. This commercial antibody also failed to discriminate between K27-, K29-, K48-, and K63-linked Ub chains in IP assays (Figure 4E). Collectively, these results establish our K27-IgG-21 (hereafter referred to as K27-IgG) as a reliable, conformation-specific tool for detecting K27-linked Ub chains.

**Figure 4.**
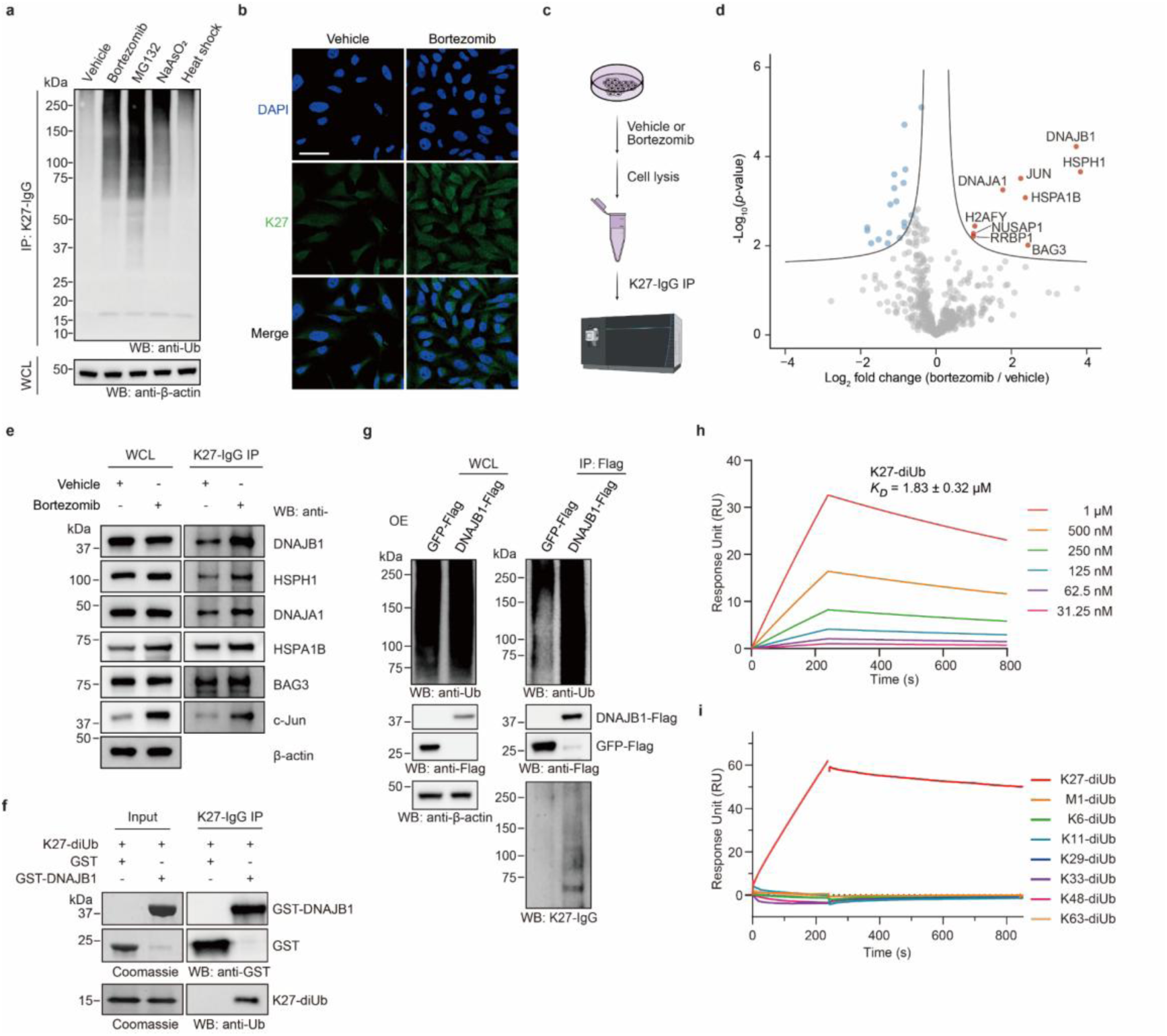
Proteomic and biochemical identification of DNAJB1 as a specific reader of K27-linked ubiquitin. **(a)** Western blot analysis of K27-liniked ubiquitination following treatment of HEK293T cells with bortezomib (1 μM, 4 h), MG132 (10 μM, 4 h), sodium arsenite (250 μM, 4 h), or heat shock (43 °C, 30 min). K27-linked ubiquitin was enriched by immunoprecipitation with K27-IgG. **(b)** Immunofluorescence imaging of K27-linked ubiquitin (green) in HeLa cells treated as in (a). Scale bar, 50 μm. **(c)** Schematic of the native K27-IgG immunoprecipitation workflow for proteomic profiling. **(d)** Volcano plot of significantly enriched proteins identified in the proteomic analysis (n = 4, biological replicates). Proteins significantly enriched (FDR < 0.05, *s_0_* = 0.1) are shown in red, and significantly depleted in blue. Key interactors of interest are highlighted. **(e)** Western blot validation of key proteomic hits enriched by K27-IgG native immunoprecipitation from bortezomib-treated HEK293T cells. HSPA1B and HSPH1 were detected using anti-HSP70 and anti-HSP105/110 antibodies, respectively. **(f)** Western blot of a pull-down assay. Purified GST-DNAJB1 or GST (as a negative control) was incubated with K27-linked diUb and subjected to enriched using K27-IgG. **(g)** Western blot of K27-linked polyUb co-IP with overexpressed DNAJB1-Flag from bortezomib-treated HEK293T cells. Cell lysates were subjected to anti-Flag immunoprecipitation, followed by Western blot with K27-IgG. **(h)** SPR analysis showing DNAJB1 binds to synthetic K27-linked diUb with a measurable *K_D_* of 1.83 ± 0.32 μM, respectively. **(i)** SPR analysis of DNAJB1 binding to all eight linkage diUb linkages (2 μM). Only K27-linked diUb showed detectable binding, indicating high binding specificity of DNAJB1. All image panels and gel panels in this figure represent at least two independent experiments.

### Proteome-wide identification of K27-polyUb interactions using K27-IgG

We next sought to detect K27-linked Ub signals in intact cellular systems. However, no detectable signals were observed via either direct western blotting of whole cell lysate or IP samples (Figure S5A). This inability to detect signals underscores the longstanding challenge of studying K27-linked ubiquitination—namely, its low endogenous abundance and rapid turnover^27,28^. When cells were treated with several cellular stressors, including bortezomib (a proteasome inhibitor; 1 μM, 4 h), MG132 (another proteasome inhibitor; 10 μM, 4 h), sodium arsenite (an oxidative stress response inducer; 250 μM, 4 h), and heat shock (43 °C, 30 min). All treatments led to a substantial increase in K27-linked Ub signals assessed by both IP-blotting (Figure 4A) and immunofluorescence (IF) (Figure 4B; Figure S5B), among which effects of proteasome inhibition by bortezomib and MG132 were the most pronounced. We therefore immunoprecipitated K27-linked Ub signals from HEK293T cells treated with bortezomib (1 μM, 4 h) (Figure 4C, Figure S5C). Proteomic profiling identified 27 differentially enriched proteins (four biological replicates; FDR < 0.05, *s_0_* = 0.1), including 9 upregulated and 18 downregulated proteins (Figure 4d; Table S2). Specifically, DNAJB1, HSPH1, JUN, HSPA1B, DNAJA1, H2AFY, RRBP1, NUSAP1 and BAG3 were significantly enriched (Figure S5D), in which JUN has previously been reported as a substrate of K27-linked ubiquitination^35^. Further gene ontology (GO) analysis of these interactors revealed strong associations with protein folding and molecular chaperones (Figure S5E).

### DNAJB1 specifically recognizes K27-linked ubiquitin

To validate our proteomic findings, we first immunoblotted K27-IgG IP eluates from bortezomib-treated HEK293T cells for six most enriched proteins: DNAJB1, HSPH1, JUN, HSPA1B, DNAJA1, and BAG3. Notably, all six proteins were enriched in the K27-IgG IP fractions, with DNAJB1 showing the most prominent enrichment (Figure 4E), consistent with the proteomic data. IF imaging of HeLa cells further revealed partial co-localization between intracellular DNAJB1 (red) and K27-linked ubiquitination (green) under bortezomib treatment, supporting the physiological relevance of their interaction (Figure S6A).

Pull-down assays confirmed that recombinant GST-DNAJB1 binds to purified K27-linked diUb *in vitro* (Figure 4F). This binding was further validated using endogenous polyUb from whole cell lysates (Figure S6B). In a complementary approach, overexpressed DNAJB1-Flag in HEK293T cells was co-immunoprecipitated with endogenous K27-linked polyUb upon bortezomib treatment (Figure 4G). We next quantified the binding affinity and specificity of this interaction. SPR analyses showed K27-diUb and tetraUb bound DNAJB1 with *K_D_* values of 1.83 ± 0.32 μM and 3.42 ± 1.58 μM, respectively (Figure 4H; Figure S6C-D). Comparative SPR of all eight diUb linkages revealed DNAJB1 exhibited measurable binding only to K27-linked diUb (Figure 4I). Together, these results establish DNAJB1 as a specific “reader” of K27-linked ubiquitination.

### K27-polyUb has a potential chaperone-like activity

Interestingly, the identified DNAJB1, HSPH1, HSPA1B, DNAJA1, and BAG3 were components of a well-established, HSPA1B-centered network known as the HSP70 molecular chaperone machinery (Figure S7A). To investigate the potential function of K27-linked ubiquitination in the context of binding with DNAJB1, we generated DNAJB1 knocked-out cell line using CRISPR-Cas9 and rescued the knockout by transfecting DNAJB1. Upon DNAJB1 knockout, we observed a modest increase in K27-linked ubiquitin signal; rescue of DNAJB1, however, led to a decrease. (Figure S5A-B). The observed results lead to the hypothesis that K27-linked ubiquitination and DNAJB1 might work through compensatory mechanisms.

To dissect the biochemical basis of this compensatory relationship, we next investigated how K27-linked Ub chains directly influence the activity of the HSP70-DNAJB1 chaperone system. For this purpose, we synthesized structurally defined tetraUb^60,61^ with K27 linkage to assess its impact on HSP70-DNAJB1-mediated chaperone activity. In the luciferase refolding assay, the luciferase activity was examined after sequential denaturation and refolding with HSPA1B, DNAJB1, K27-tetraUb or their combinations to evaluate the chaperone-like activity of K27-linked ubiquitination. We found that K27-linked tetraUb significantly enhanced HSPA1B-mediated luciferase refolding. More interestingly, K27-linked tetraUb alone can facilitate luciferase refolding in the absence of HSPA1B (Figure 5C).

**Figure 5.**
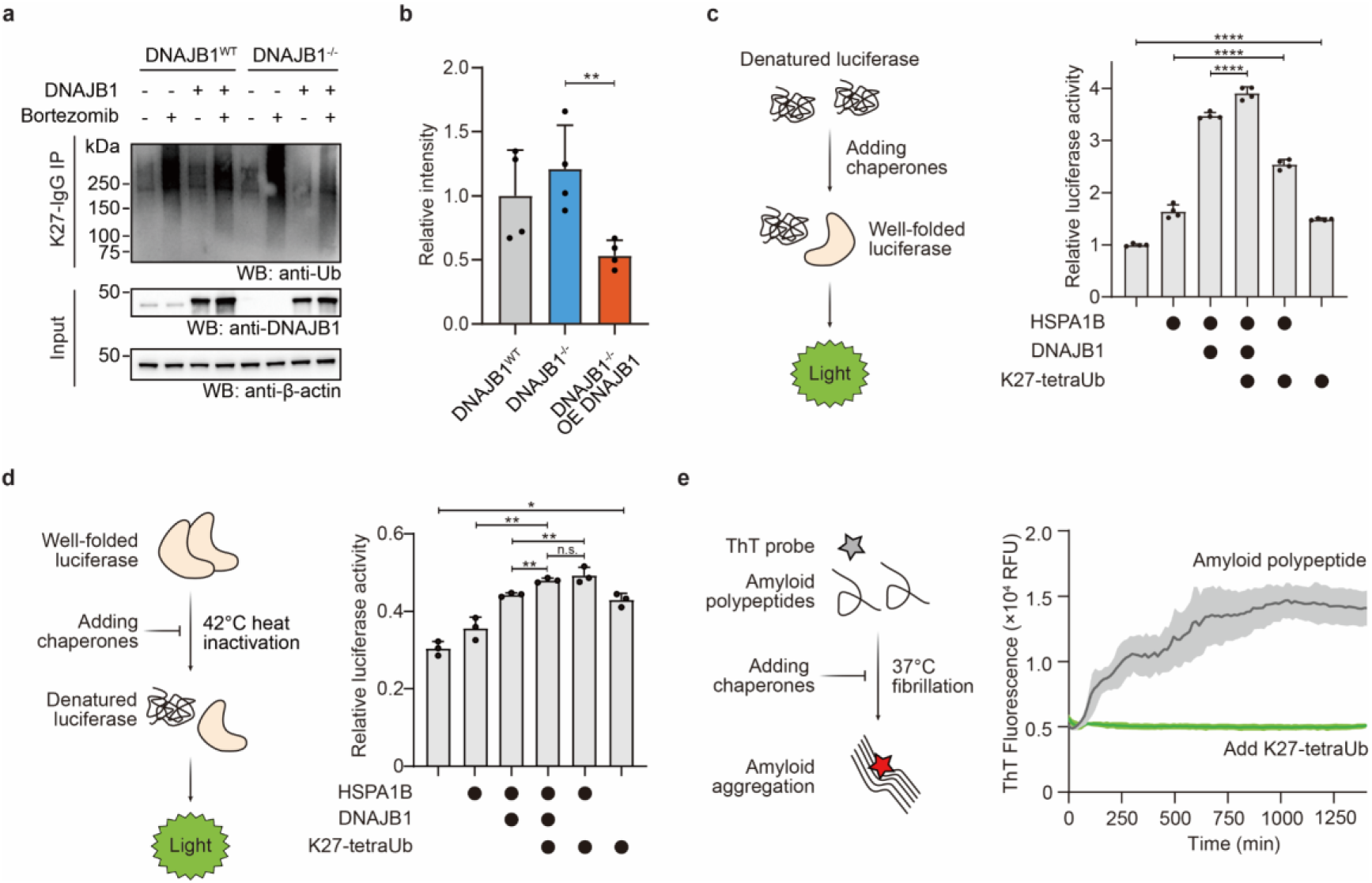
K27-linked ubiquitin modulates the HSP70 chaperone system and potentially suppresses protein aggregation. **(a-b)** Western blot detection (a) and quantification (b) of endogenous K27-linked ubiquitin levels in DNAJB1-knockout and DNAJB1-overexpressing HEK293T cells (n = 4, biological replicates; one-tailed Student’s t-test; **, p < 0.01). **(c)** Endpoint quantification (2 h) of luciferase refolding assays containing HSPA1B, DNAJB1, K27-linked tetraubiquitin and their combinations, showing that K27-linked ubiquitin enhances HSP70 activity and also independently promotes protein refolding. Data are presented relative to the initial value. (n = 4; one-way ANOVA with Tukey’s multiple comparisons test; ****, p< 0.0001) **(d)** Endpoint quantification (6 min) of luciferase inactivation assays showing that K27-linked tetraUb (1 μM) synergizes with the HSP70-DNAJB1 complex to prevent thermal inactivation. K27-linked ubiquitin alone also delays inactivation, indicating a chaperone-independent protective effect. Data are presented relative to the initial value (n = 3; one-way ANOVA with Tukey’s multiple comparisons test; n.s., not significant; *, p < 0.05; **, p <0.01). **(e)** *In vitro* fibrillization assays monitored by ThT fluorescence in the presence of K27-linked tetraUb, showing that K27-linked tetraUb alone robustly suppresses tau fibril formation, supporting a direct anti-aggregation role. All image panels and gel panels in this figure represent at least three independent experiments.

In a complementary luciferase inactivation assay, luciferase activity was examined after thermal inactivation in the presence of HSPA1B, DNAJB1, K27-tetraUb or their combinations. Again, we observed K27-linked Ub chain synergized with HSP70 to prevent thermal inactivation of luciferase, with an effect comparable to DNAJB1 itself. Additionally, K27-linked tetraUb alone delayed inactivation, demonstrating a direct, chaperone-independent protective role (Figure 5D; Figure S7B). These observations suggest that K27-linked ubiquitination replenishes some of the chaperone function.

To validate this hypothesis, we performed fibrillization assays with human islet amyloid polypeptide (hIAPP). K27-linked tetraUb was observed to potently suppress fibril formation, revealing intrinsic anti-aggregation activity specific to this Ub chain topology (Figure 5E). Taken together, these findings indicate a chaperone-like role of K27-linked Ub chains: on one hand, they can enhance HSP70-mediated luciferase refolding and protect against thermal denaturation; on the other hand, K27-linked ubiquitin can directly facilitate luciferase renaturation, delay inactivation, and inhibit hIAPP aggregation independently of chaperones.

### Identification of UBE2Q1 generating K27-linked di-Ubiquitin

Our above-described IP protocol is based on a native lysis environment for ubiquitinated sample preparation and enrichment process, which may lead to insufficient protein extraction (membrane related or condensation related), unstable ubiquitin signal (trimmed by deubiquitinating and proteasomes). To profile the substrates of K27-linked ubiquitination, we adopted the recently reported Denatured-Refolded Ubiquitinated Sample Preparation (DRUSP) method, combined with K27-IgG-based IP-MS. This strategy could break the non-covalent protein interactions while further enhance the detection of low-abundant ubiquitination events (Figure 6A)^62,63^. Using the DRUSP method, we achieved significantly enhanced detection of K27-linked Ub signals in bortezomib-treated HEK293T cell lysates (Figure 6B; Figure S8A-B). Proteomics profiling of K27-linked ubiquitination in these cells identified 173 enriched proteins (n = 4, biological replicates; FDR < 0.05, *s_0_* = 0.1) (Figure S8C; Table S3). These enriched proteins are involved in Wnt signaling (SIAH1 and CTNNB1), mitochondrion (MRPS28, PDF, ISOC2, CTNNB1, and NDUFS4) and neurodegeneration (AP2S1, CTNNB1, and NDUFS4) (Figure S8D). Notably, most HSP70 family members, including DNAJB1, showed no significant enrichment in this DRUSP-based IP-MS dataset (Figure S8E-F). This result confirms that the denaturing lysis buffer effectively eliminated non-covalent protein-protein interactions, supporting the identification of high-confidence K27-ubiquitinated substrates.

**Figure 6.**
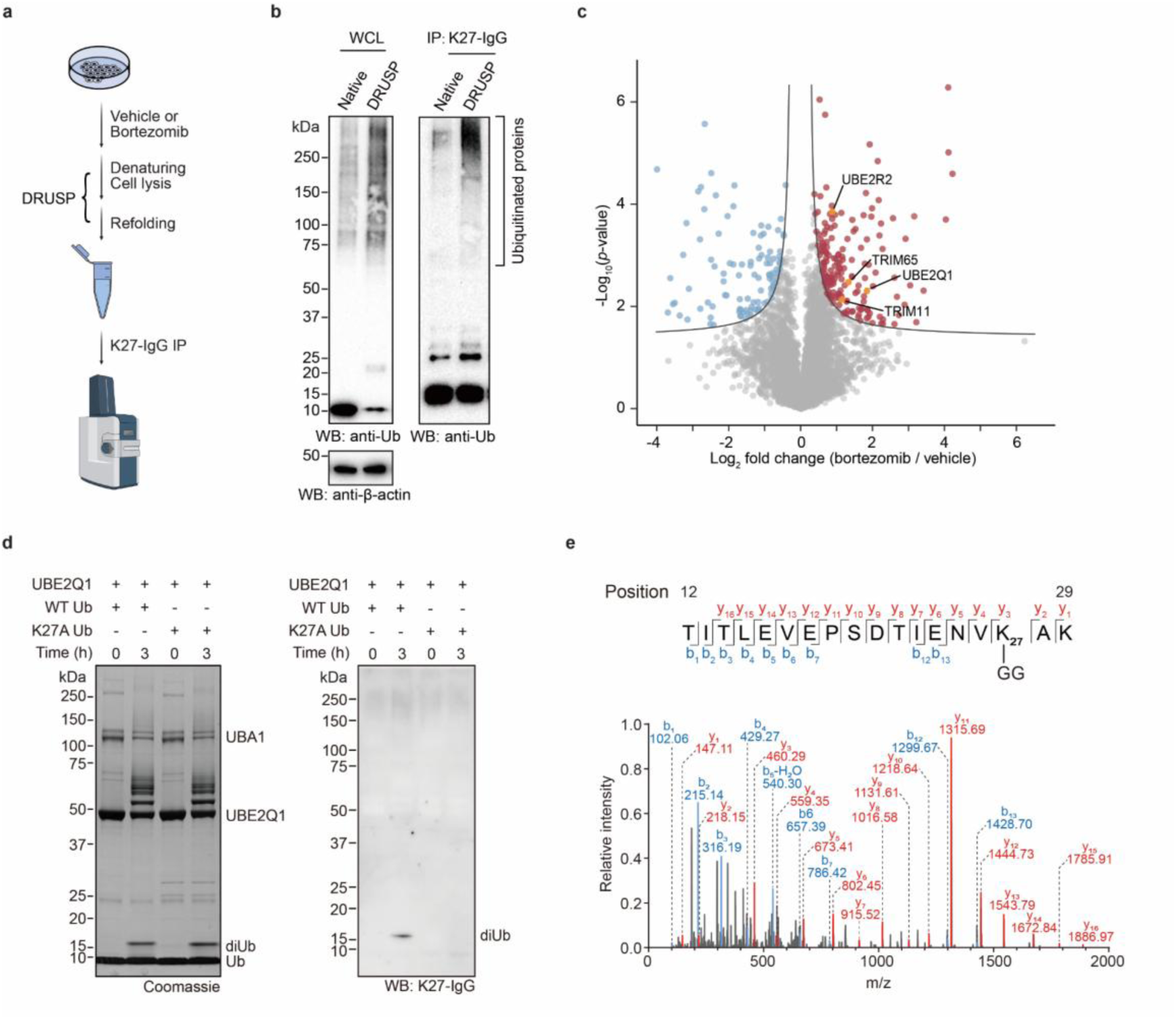
Denaturing proteomic profiling identifies UBE2Q1 as a potential writer of K27-linked ubiquitin. **(a)** Schematic overview of DRUSP-based K27-IgG IP and subsequent proteomic profiling. **(b)** Comparison of enrichment efficiency for K27-linked ubiquitinated substrates between native IP and DRUSP-based IP. **(c)** Volcano plot of differentially enriched proteins identified by proteomic analysis (n = 4, biological replicates). Significantly enriched proteins (FDR < 0.05, *s_0_* = 0.1) are shown in red, and depleted proteins are in blue. The E2 and E3 enzymes, including UBE2Q1, are indicated in orange. **(d)** *In vitro* ubiquitination assay showing that recombinant UBE2Q1 catalyzes the formation of K27-linked diUb, as detected by Western blot with K27-IgG. **(e)** Tandem mass spectrum confirming the presence of K27-linked diglycine (GG) peptides derived from UBE2Q1-assembled diUb. All gel panels in this figure are representative at least three independent experiments.

Intriguingly, among the most significantly enriched proteins, we identified several E2 ubiquitin-conjugating enzymes and E3 ubiquitin ligase (Figure 6C). To investigate potential “writers” of K27-linked ubiquitination, we expressed and purified recombinant full-length UBE2Q1, UBE2R2, and TRIM11, TRIM65, and then performed *in vitro* ubiquitination assays (Figure S9A). Western blot analysis of reaction products using K27-IgG revealed that UBE2Q1 catalyzes formation of K27-linked diUb (Figure S9B). Critically, this activity was abolished by K27A mutation of ubiquitin (Ub-K27A), confirming that the diUb formation depends on Ub’s K27 residue (Figure 6D). Furthermore, the MS/MS spectrum confirmed K27-linked isopeptide formation in UBE2Q1-assembled diUb (Figure 6E; Table S4). These results establish UBE2Q1 as an E2 enzyme that directly assembles K27-linked ubiquitin chains.

## Discussion

K27-linked ubiquitination remains one of the most challenging ubiquitin modifications to study in depth, primarily due to the lack of specific tools capable of recognizing its native structure for its enrichment. Here, we developed conformation-specific antibodies targeting the unique architecture of K27-linked Ub chains by integrating protein synthetic chemistry and phage display screening. High-resolution co-crystal structures further elucidated the molecular basis for their selective recognition of the K27-diUb interface. Upon reformatting into a full-length human IgG, the optimized antibody (K27-IgG) achieved nanomolar affinity (*K_D_* = 4.66 nM) and enabled sensitive IF detection and IP of endogenous K27-linked Ub signals in cellular systems.

Leveraging this tool, we implemented complementary proteomic approaches to map the cellular K27-linked Ub landscape in HEK293T cells under proteotoxic stress. Native IP-MS identified molecular chaperone DNAJB1 as a direct interactor of K27 Ub chains, linking this modification to chaperone-mediated proteostasis. Denaturing IP-MS further revealed UBE2Q1 as an E2 enzyme responsible for K27 linkage assembly. Collectively, these findings establish K27-IgG as a transformative reagent for deciphering the molecular functions of K27-linked ubiquitination.

Critically, DNAJB1 (an HSP70 co-chaperone) is reported to actively prevent the aggregation of nascent or stress-denatured polypeptides and aids HSP70 in promoting the proper folding, refolding, or targeted degradation of client proteins^64–70^. The interaction between K27-linked Ub chains and DNAJB1 thereby connects this atypical ubiquitination to a chaperone-mediated protein quality control (PQC) function. We functionally validated this mechanism by reconstituting the chaperone-like activity of K27-linked ubiquitination *in vitro* using luciferase-refolding and amyloid fibril formation assays. These discoveries provide a unifying mechanistic framework for previously disparate observations regarding K27-linked ubiquitination. For example, PTEN modification by K27 ubiquitination was reported to affect its stability and subcellular localization^22^, a phenomenon that could be interpreted as K27 Ub chains facilitating chaperone engagement to prevent dimerization. K27 ubiquitination by WSB1 was proposed to drive LRRK2 aggregation^16^, which may represent a protective sequestration process coordinated by the DNAJB1-HSP70 axis. Similarly, MARCH8-mediated K27 ubiquitination promotes receptor degradation^37^, consistent with a role for chaperone-assisted extracting of membrane substrates toward degradation pathways. In addition, previous studies showed that multiple stress-inducible E3 ligases, including Parkin^32^, Rhbdd3^20^, NEDD4^31^, HACE1^33,34^, TRIM23^39^, RNF168^40^, LTN1^71^, and CHIP^72^ and assemble K27 linkages under proteotoxic stress, supporting its role in sustained adaptive responses. Collectively, we propose that K27 chains could represent a higher-order “chaperone code”. The precise molecular mechanisms remain to be clarified.

### Limitations of the study

Despite the identification of DNAJB1 and UBE2Q1 as a reader and writer, respectively, of K27-linked Ub chains, structural insights into the DNAJB1-K27 interaction and UBE2Q1-mediated Ub chain assembly remain lacking. Further studies are needed to elucidate these mechanisms.

## Resource availability

### Lead contact

Further information and requests for the resources and reagents may be directed to the corresponding author, Yuanyuan Yu.

## Data availability

The atomic model of K27-linked diUb in complex with sAB-K27 has been deposited in the Protein Data Bank (PDB) under the accession code 9WYW. The MS proteomics data have been deposited to the ProteomeXchange Consortium (https://proteomecentral.proteomexchange.org) via the iProX partner repository^85,86^ with the dataset identifier PXD068933. Source data are provided with this paper.

## Acknowledgements

This work was supported by the National Key R&D Program of China (2023YFC2306500 to Y.Y.; 2022YFC3401500 to L.L.; 2023YFA0915300 to M.P.), and the National Institutes of Health under grant number R35GM143052 to M.Z.. We thank the National Natural Science Foundation of China (22207070, 22477076 to Y.Y.; 92253302, T2488301, 22137005, and 22227810 to L.L.; 22277073 and 92253302 to M.P.; 22407086 to Y.W.; 22407085 to Q.Z.; 22307071 to Q.Q.) for the support. We thank the Foundation of the National Facility for Translational Medicine (Shanghai) (TMSK-2024-110 to Y.Y.), New Cornerstone Science Foundation (to L.L.), Shanghai Pilot Program for Basic Research - Shanghai Jiao Tong University (21TQ1400224 to M.P.), Foundation of Muyuan Laboratory (118602240 to M.P.), Fundamental Research Funds for the Central University (to M.P.), Shanghai Jiao Tong University 2030 Initiative (WH510363003/003 to M.P.). Chenguang Program of Shanghai Education Development Foundation and Shanghai Municipal Education Commission (23CGA14 to Q.Z.). Structural characterization is based upon research conducted at the Northeastern Collaborative Access Team beamlines, which are funded by the National Institute of General Medical Sciences from the National Institutes of Health (P30 GM124165). The Eiger 16M detector on the 24-ID-E beam line is funded by a NIH-ORIP HEI grant (S10OD021527). This research used beamtime awards from the Advanced Photon Source, a U.S. Department of Energy (DOE) Office of Science User Facility operated for the DOE Office of Science by Argonne National Laboratory under Contract No. DE-AC02-06CH11357.

## Author Contributions

Y. Yu, C. Han, L. Liu, M. Zhao and M. Pan proposed the idea, designed the experiments and interpreted the results. Q. Qu, Q. Zheng and Y. Han synthesized all diUb molecules. S.K. Erramilli and A.A. Kossiakoff performed sAB selection and C. Han, Z. Su, Y. Duan and X. Zhai conducted biochemical evaluation. Y. Yu, M. Zhao and J. Li conducted crystallographic screening and structure determination. C. Han, Y. Weng and Q. Zheng performed proteomic analysis. C. Han and Y. Duan carried out cell-based experiments. Y. Yu, C. Han, M. Pan, Q. Zheng, Y. Weng, Q. Qu, L. Liu read, discussed and analyzed the manuscript. Y. Yu, L. Liu and M. Pan supervised the project.

## Methods

### Cell culture and transfection

HEK293T cells (CRL-3216) were obtained from the American Type Culture Collection (ATCC). HeLa cells (TCHu187) and U-2OS cells (TCHu88) were kindly provided by Cell Bank, Chinese Academy of Sciences. HEK-293T, HeLa, and U-2OS cells were cultured at 37 °C and 5% CO_2_ in Dulbecco’s Modified Eagle’s Medium (Gibco, 11995500) supplemented with 10% Fetal Bovine Serum (PAN Biotech, ST-3302), 100 units/mL of penicillin, and 100 μg/mL of streptomycin (Gibco, 15140122). HEK-293F cells were cultured at 37 °C, 120 rpm and 5 %CO_2_ in SMM 293-TII Expression medium (Sino Biological, M293TII). Cells were routinely confirmed to be mycoplasma-free. Adherent cell lines were transfected using Lipofectamine 3000 (Invitrogen, L3000015). HEK-293F cells were transfected using PEI MAX 40K (Polysciences, 24765). All transfections were performed according to the respective manufacturer’s instructions.

### Plasmid constructs

sABs were cloned into the *HindIII*/*SalI* sites of the expression vector pRH2.2. IgGs were derived from sABs, the light chain was cloned into *NotI*/*BamHI* site and the heavy chain was cloned into *NotI*/*NheI* sites of PBR322-based human Ig-λ expression vector, as previously described^73^. Human DNAJB1 (NM_006145.3) was obtained from MiaoLing Plasmid Platform and subcloned into the *BamHI*/*XhoI* sites of pGEX-6p-1 for bacterial expression or into the same sites of pcDNA3.1(+) with a C-terminal DYKDDDDK tag for mammalian overexpression. Human HSPA1B (NM_005346.6) was obtained from MiaoLing Plasmid Platform and subcloned into the *BamhI*/*XhoI* sites of the pET-28a(+) vector, which contains an N-terminal His_6_ tag and a Ulp1 cleavage site. Human UBE2K (NM_005339.5) was synthesized by GenScript and cloned into the *NheI*/*XhoI* sites of pET-28a(+) for bacterial expression. Human UBE2Q1 (NM_017582.7), UBE2R2 (NM_017811.4), UBCH5C (NM_181891.3) and UBE2G1 (NM_003342.5) was obtained from MiaoLing Plasmid and cloned into the *NcoI*/*NotI* sites of pET-28a(+) with N-terminal His_6_ tag for bacterial expression. Human TRIM11 (NM_145214.3) and TRIM65 (NM_173547.4) was obtained from MiaoLing Plasmid and cloned into the *BamHI*/*XhoI* sites of pGEX-6p-1 for bacterial expression. For CRISPR-Cas9 genome knockout, double strands of single guide RNA targeting hDNAJB1 (5’-CACCGCGGGTCGCTGAGCACGTCGT-3’, and 5’-AAACACGACGTGCTCAGCGACCCGC-3’) were syntheses by Sangon Biotech and cloned into the *BbsI* site of PX459 backbone (Addgene #62988; obtained from MiaoLing Plasmid), termed PX459-sgDNAJB1#2.

### Antibodies

Antibodies used in this study for immunofluorescence (IF) or western blot (WB) were listed below. Primary antibodies: sAB-K27-9 (this study, WB 0.4 μg/mL); sAB-K27-21 (this study, WB 0.4 μg/mL); K27-IgG-9 (this study, WB 0.4 μg/mL); K27-IgG-21 (this study, WB 0.4 μg/mL, IF 2.5 μg/mL); K29-IgG (previous study^15^, WB 0.2 μg/mL); Anti-ubiquitin (Abcam, ab19247, WB 1:5000); Anti-K48-ubiquitin (Abcam, ab140601, WB 1:2000); Anti-K63-ubiquitin (Abcam, ab179434, WB 1:2000); Anti-K27-ubiquitin (Abcam, ab181537, WB 1:2000); Anti-DNAJB1 (Proteintech, 13174-1-AP, WB 1:5000, IF 1:400); Anti-DNAJA1 (Proteintech, 11713-1-AP, WB 1:5000); Anti-HSP70 (Proteintech, 10995-1-AP, WB 1:5000); Anti-HSP105/HSP110 (Abcam, ab109624; WB 1:2000); Anti-c-Jun (CST, 9165, WB 1:2000); Anti-Bag3 (Proteintech, 10599-1-AP, WB 1:5000); Anti-β-actin (CST, 8457, WB 1:10000), Anti-DYKDDDDK Tag (CST, 14793, WB 1:5000), Anti-GST tag (CST, 2622, WB 1:5000). Secondary antibodies: Peroxidase-conjugated anti-rabbit IgG (CST, 7074, WB 1:5000); Peroxidase-conjugated anti-mouse IgG (CST, 7076, WB 1:5000); Peroxidase-conjugated goat anti-human IgG (F(ab)_2_ fragment specific) (Jackson, 109-036-006, WB 0.2µg/mL); Peroxidase-conjugated donkey anti-human IgG (H+L) (Jackson, 709-035-149, WB 0.2µg/mL); Alexa Fluor 647-conjugated Goat Anti-Rabbit IgG (H+L) (Jackson, 111-605-144, IF 2 µg/mL); Alexa Fluor 488-conjugated Donkey Anti-Human IgG (H+L) (Jackson, 709-545-149, IF 2 µg/mL).

### Synthesis of tagged K27-linked diUb

Tagged K27-linked diUb was synthesized using hydrazide-based native chemical ligation (NCL) and microwave-assisted solid phase peptide synthesis (MW-SPPS) as described previous (Extended Data Fig. 1a)^30,54,55^. Briefly, biotinylated K27-linked diUb was divided into three peptide segments, namely fragment Ub[Met_1_-Phe_45_]-NHNH_2_, fragment Ub[Cys_46_-Gly_76_]-NH_2_, and fragment Ub[Cys_46_(Acm) -Gly_75_]-Gly-[Met_1_-Lys_27_-Phe_45_]-NHNH_2_. Ala_46_ of each Ub subunit was temporarily mutated to Cys for NCL. The isopeptide bond at Lys_27_ of fragment Ub[Cys_46_(Acm)-Gly_75_]-Gly-[Met_1_-Lys_27_-Phe_45_]-NHNH_2_ was constructed using the orthogonal building block Fmoc-Lys(Alloc)-OH rather than normal Fomc-Lys(Boc)-OH. An acetamidomethyl (Acm) group was used to protect the thiol group of N-terminal Cys of to avoid oligomerization or self-cyclization. All the peptides were synthesized using standard Fmoc SPPS protocols under microwave conditions (CEM Liberty Blue). The insertion of PEG linker was constructed using the building block Fmoc-mini-PEG, and biotin was directly condensed to the N termini. Fragment Ub[Cys(Acm)_46_-Gly_75_]-Gly-[Met_1_-Lys_27_-Gly_76_]-NH_2_ was synthesized by hydrazide-based native chemical ligation of fragment Ub[Cys_46_(Acm)-Gly_75_]-Gly-[Met_1_-Lys_27_-Phe_45_]-NHNH_2_ with Ub[Cys_46_-Gly_76_]-NH_2_ followed by Acm deprotection. Finally, fragment Ub[Met_1_-Phe_45_]-NHNH_2_ and fragment Ub[Cys_46_(Acm)-Gly_75_]-Gly-[Met_1_-Lys_27_-Gly_76_]-NH_2_ were ligated to generate the full-length diUb, which was subjected to desulfurization to convert Cys_46_ in the two Ub units into Ala_46_.

### Phage display selection

Phage display was carried out as previously established protocols with minor modifications^15,56,57^. Briefly, four rounds of biopanning were performed. In the first round, 300 nM of biotinylated K27-linked diUb was immobilized on 250 μL of Streptavidin Paramagnetic Particles (Promega, Z5481) and incubated with Fab Library E (kind gift from A. Koide) for 1 h at room temperature. Beads were pre-equilibrated and washed using selection buffer (20 mM HEPES-NaOH, pH 7.5, 150 mM NaCl, 0.5 % bovine serum albumin, 0.05 % Tween-20). Bound phages were used to infect log-phase *E. coli* XL1-Blue, and amplified overnight in 2×YT medium supplemented with M13 helper phage. To increase selection stringency, 1.5 μM mono-ubiquitin was included as a competitor in rounds two to four, and the concentration of K27-linked diUb was sequentially reduced to 150, 75, and 15 nM, respectively. Phages enriched from each round were directly used as input for the subsequent round. The first round was performed manually, and subsequent rounds were performed semi-automatically using a Kingfisher magnetic beads handler.

### Purification of synthetic antibody fragment

Plasmids encoding sAB-K27s were transformed to *E. coli* BL21(DE3) (TransGen Biotech, CD601). Protein expression was induced with 0.5 mM isopropyl β-D-thiogalactoside (IPTG) for 4 h at 37 °C. Cells were pelleted by centrifugation at 4,000 × *g* for 20 min at 4 °C and resuspended in lysis buffer (20 mM Tris-HCl, pH 7.4, 500 mM KCl, 10 % glycerol and 1 mM phenylmethylsulfonyl fluoride (PMSF)). The cell suspension was homogenized, incubated at 65 °C for 30 min and then clarified by centrifugation at 40,000 × *g* for 1 h. The supernatant was loaded onto a MabCap At 4FF column (Smart-Lifescience, SA023) pre-equilibrated with lysis buffer, and processed using an automated program on ÄKTA Pure FPLC system (Cytiva). The sABs were eluted with 0.1 M glycine, pH 3.0, followed by cation-exchanged chromatography using a SOURCE 15S 10/200 GL (Cytiva, 11001288). The proteins were eluted using a linear NaCl gradient by mixing buffer A (50 mM sodium acetate, pH 4.5) with buffer B (50 mM sodium acetate, pH 4.5, 1 M NaCl). Purified sABs were dialyzed overnight against ice-cold phosphate-buffered saline (Gibco, 10010023).

### Characterization of binding affinity

SPR experiments were performed on a Biacore 8K+ instrument (Cytiva) using an NTA sensor chip (Cytiva, BR100407) and HBS-T running buffer (10 mM HEPES-NaOH, pH 7.4, 150mM NaCl, 50 μM ethylene diamine tetraacetic acid (EDTA), 0.005% Tween-20). The sensor surface was activated with 0.5 mM NiCl_2_ for 60 s, followed by immobilization of His_6_-K27 diUb at 10 μL/min for 60 s in experimental channels only. After optimizing ligand immobilization based on response units (RU), serial dilutions of sAB-K27s or K27-IgGs were injected at 30 μL/min for 120 s over both experimental and reference channels. Regeneration was performed with either 350 mM EDTA or 50 mM NaOH. Data were fitted using a 1:1 Langmuir binding model in Biacore Insight Evaluation software v5.0.18 (Cytiva). All experiments were performed in triplicate independent replicates.

### Purification of K27- IgG

Plasmids encoding the heavy and light chain of K27-IgG were co-transfected into HEK293F cells at a 1:2 ratio for transient overexpression. After 5-7 days of culture, cells were removed by centrifugation at 1,000 × *g* for 10 minutes, and the clarified supernatant was collected. The supernatant was loaded onto a MabCap At 4FF column (Smart-Lifescience, SA023) pre-equilibrated with PBS. The K27-IgG was eluted with Elution Buffer (25 mM H_3_PO_4_, pH 2.8, 100 mM NaCl) and immediately neutralized with Neutralization Buffer (0.5 M Na_3_PO_4_, pH 8.0). The eluted K27-IgG was dialyzed overnight against ice-cold PBS.

### Purification of proteins used for *in vitro* ubiquitination assay

Recombinant proteins were expressed in *E. coli* BL21(DE3) grown in Luria-Bertani (LB) medium containing the appropriate antibiotic (100 µg/mL ampicillin or 50 µg/mL kanamycin). At an optical density at 600 nm (OD_600_) of 0.8, protein expression was induced with 0.5 mM IPTG, and cultures were incubated for 16 hours at 18°C. Cells were harvested by centrifugation (4,000 × *g*, 20 min, 4 °C) and resuspended in lysis buffer (20 mM Tris-HCl, pH 8.0, 500 mM KCl, 10 % glycerol, 5 mM MgCl_2_, 1 mM Tris(2-carboxyethyl)phosphine hydrochloride (TCEP), 1 mM PMSF). Following lysis by a high-pressure homogenizer, the lysate was clarified by centrifugation at 17,000 rpm for 90 min at 4 °C. The clarified supernatant was loaded onto affinity chromatography using either Ni-NTA or Glutathione Beads. Beads were washed with Buffer (20 mM Tris-HCl, pH 8.0, 150 mM NaCl) supplemented with 20 mM imidazole for the Ni-NTA beads. Bound proteins were eluted with Buffer supplemented with either 400 mM imidazole for the Ni-NTA beads or 10 mM reduced glutathione for glutathione beads. Eluted proteins were dialyzed overnight at 4 ℃ in the dialysis buffer (20 mM Tris-HCl, pH 8.0, 150 mM NaCl and 1 mM TCEP). Finally, protein was purified by size exclusion chromatography on a Superdex 75 or 200 Increase 10/300 GL column (Cytiva, 29148721 or 28990944) pre-equilibrated with the dialysis buffer.

### Generation of polyubiquitin chains

Synthesis of ubiquitin chains *in vitro* was performed as previously described^74^. All reactions contained 1 mM wild-type ubiquitin, 1 µM UBA1, and a reaction buffer (50 mM HEPES-NaOH, pH 7.5, 150 mM NaCl, 10 mM ATP, 10 mM MgCl_2_, 0.6 mM dithiothreitol (DTT)). K29-linked ubiquitin chains were generated by the addition of 20 µM UBCH7 and 20 µM UBE3C, K48 with 20 µM UBE2K, and K63 with 20 µM Ubc13 and 20 µM Mms2. Reactions were incubated at 37 °C for 4 h (UBE3C, UBE2K) or 12 h (Ubc13/Mms2). The reactions were monitored by Coomassie staining and western blot.

### Crystallization of sAB-K27 and K27-linked diUb complex

To form the complex, sAB-K27 and K27-linked diUb were mixed at a 1:1.5 molar ratio, following by isolation of the complex using size exclusion chromatography on a Superdex 75 Increase 10/300 GL column (Cytiva, 29148721) equilibrated in a buffer (20 mM Tris-HCl, pH 7.4, 150 mM NaCl). The sAB-K27-diubiquitin complex was confirmed by Coomassie staining. crystals containing SAB-K27-9 and K27-diUb grew in 1 weeks at 21 °C in hanging drops from a 1:1 mixture of protein (11.8 mg/mL in 20 mM Tris-HCl, pH 7.4, 150 mM NaCl) and well solution (0.005 M cobalt(II) chloride hexahydrate, 0.005 M Nickel(II) chloride hexahydrate, 0.005 M Cadmium chloride hydrate, 0.005 M Magnesium chloride hexahydrate, 0.1 M HEPES, pH 7.5, 12 % w/v PEG 3,350) (F4, Index, Hampton Research). Crystals containing SAB-K27-21 and K27-diUb were obtained under similar conditions with protein at 15.1 mg/mL, using well solution (0.1 M HEPES, pH6.5, 10 % (w/v) PEG 6000, final pH 7.0) (C4, JCSG plus, Hampton Research). Diffraction-quality crystals were flash-frozen in liquid nitrogen with 23 % glycerol as cryo-protectant.

### Data collection and structure determination

The crystal structure was determined following previously established crystallographic workflow^15^. Briefly, X-ray diffraction data were collected at the Northeastern Collaborative Access Team beamline 24-ID-E at the Advanced Photon Source. The datasets were subsequently processed and scaled using XDS/XSCALE^75^. Structures were solved by molecular replacement using PHASER (CCP4) and refined with REFMAC5 (CCP4)^76,77^. the humanized 4D5 Fab (PDB 1FVE) and ubiquitin (PDB 1UBQ) were used as search models. Final data collection and refinement statistics are summarized in Supplementary Table 1. All structure figures were prepared using ChimeraX 1.8^78^ (University of California, San Francisco) or open-source PyMOL 3.2.0a (Schrödinger).

### Western blot

Prior to electrophoresis, protein samples were denatured by boiling for 10 min in Laemmli Sample Buffer (Bio-Rad, 1610747). The samples were then separated on 4-20 % bis-tris gradient gels (GenScript, M00657) and subsequently transferred onto a polyvinylidene difluoride (PVDF) membrane (Bio-Rad, 1620177). After transfer, membranes were blocked for 1.5 h at room temperature in 5 % (w/v) skim milk (BD, 232100) in TBS-T buffer (20 mM Tris-HCl, pH 7.4, 150 mM NaCl, 0.05 % Tween-20). The membranes were then probed with primary antibodies overnight at 4 °C, followed by incubation with appropriate HRP-conjugated secondary antibodies for 1 h at room temperature. The blots were exposed by a ChemiDoc MP (Bio-Rad) using Clarity Western ECL Substrate (Bio-Rad, 1705060), and signals were analyzed by Image Lab Software ver6.1 (Bio-Rad)

### Immunofluorescence and microscopy

Cells were grown on Poly-D-Lysine pre-coated coverslips (WHB Scientific, WHB-24-CS-LC) and treated or transfected for 24 h. For staining, cells were fixed with 4 % paraformaldehyde (Beyotime, P0099) for 15 min, and permeabilized with 0.25 % Triton X-100 in PBS for 10 min. After blocking for 1 h with blocking buffer (Beyotime, P0260), cells were incubated with primary antibodies for 2 h, followed by incubation with fluorescently-labeled secondary antibodies for 1 h. All staining steps were performed at room temperature. Coverslips were mounted onto glass slides using DAPI Fluoromount-G (Southern Biotech, 0100-20). Images were acquired with Leica STELLARIS 5 Confocal Microscope Platforms using a 63× oil-immersion objective and processed with LAS X Office 1.4.6 software (Leica Microsystems).

### Cellular stress assays

To induce various cellular stress responses, cells were subjected to one of the following treatments. For chemical-induced stress, cells were treated for 4 h with either bortezomib (1 µM; MCE, HY-10227), MG132 (10 µM; MCE, HY-13259), or sodium arsenite (250 µM; Sigma-Aldrich, S7400). For heat-induced stress, cells were incubated at 43 °C for 30 min. Following treatment, cells were harvested for subsequent analysis of K27-linked ubiquitin by immunoprecipitation or immunofluorescence.

### GST pull-down assay

Glutathione S-Transferase (GST) pull-down assays were carried out following standard procedures. Briefly, 0.771 nmol of GST or GST-DNAJB1 was immobilized on 50 μl of GST magnetic agarose beads (Beyotime, P2258). Then, the beads were incubated with 50 μg of K27-linked diUb in HBS buffer (20 mM HEPES-NaOH, pH 7.4, 150 mM NaCl) overnight at 4 °C with gentle rotation. The beads were washed four times with HBS buffer containing 1 % (w/v) NP-40. Bound proteins were eluted with Laemmli sample buffer and analyzed by western blot.

### Cell lysis and anti-Flag immunoprecipitation

Cells were lysed by mild sonication in NETN lysis buffer (20 mM Tris-HCl, pH 8.0, 100 mM NaCl, 0.5 mM EDTA, 0.5 % (v/v) NP-40) supplemented with protease inhibitor cocktail (Roche, 4693132001) and 1 mM PMSF. The cell lysate was centrifuged at 20,000 × *g* for 30 min at 4 °C and the supernatant was collected for immunoprecipitation. Protein concentrations were determined using the BCA kit (Epizyme, ZJ102L). Anti-Flag immunoprecipitation was performed according to manufacturer’s instruction. Briefly, 5 mg of cell lysate supernatant was incubated with 40 μL of anti-Flag M2 affinity gel (Millipore, A2220) in HBS buffer overnight at 4 °C with gentle rotation. The beads were washed four times with NETN buffer and bound proteins were eluted with Laemmli sample buffer.

### Immunoprecipitation for mass spectrometry

For native immunoprecipitation, 20 µg of K27-IgG was incubated with 100 µL of pre-washed protein A+G agarose beads (Beyotime, P2055) for 4 h at 4 °C with gentle rotation. The antibody-conjugated beads were washed four times with NETN buffer and incubated with 5 mg of cell lysate supernatant overnight at 4 °C with gentle rotation. Beads were then washed three times with RIPA buffer (50 mM Tris-HCl pH 7.4, 150 mM NaCl, 1 % Triton X-100 and 0.1 % SDS). Bound proteins were eluted with 0.1 M glycine, pH 3.0, and immediately neutralized with 1 M Tris-HCl, pH 7.4. For denaturing immunoprecipitation, a previously established protocol was followed with a slight modification^63^. Briefly, cells were lysed by mild sonication in denaturing urea lysis buffer (8 M Urea, 50 mM Tris-HCl, pH 8.0, 150 mM NaCl, 0.2 % NP-40, 1 % Triton X-100, 1 % SDS, 1 % sodium deoxycholate, 10 % glycerol, 1 mM PMSF). The lysates were cleared by centrifugation at 21,000 × *g* for 30 min at 4 °C. The supernatant was reduced and alkylated with 10 mM TCEP and 40 mM 2-chloroacetamide (MCE, HY-W010629) for 15 min at room temperature. Refolding was performed by sequential buffer exchange using 10 kDa ultrafiltration tubes (Millipore, UFC9010) at 4°C. Samples were gradually exchanged with buffers (50 mM Tris-HCl, pH 7.5, 150 mM NaCl, 0.2 % NP-40, and 10 % glycerol) containing decreasing urea concentrations (8 M, 4 M, 2 M, 0) until the final urea concentration was below 0.1 M. Refolded proteins were then subjected to immunoprecipitation using K27-IgG as described above.

Eluted protein fractions were treated with 10 mM TCEP and 50 mM 2-chloroacetamide at 37 °C for 1 h with shaking. followed by 50 mM DTT. Proteins were then digested overnight at 37 °C with 20 ng/μL trypsin (Promega, V5111) and 4 ng/μL lysyl endopeptidase (Lys-C; Wako, 121-05063) with shaking. The resulting peptides were desalted using home-made StageTips with C18 membrane (CDS Empore, 2315) before LC-MS/MS analysis.

### Mass spectrometry data collection

For K27-IgG native IP sample, tryptic peptides were separated by an Aurora Elite column (75 μm i.d. × 15 cm, 1.7 μm C18) with a binary buffer system of 0.1 % formic acid in water (buffer A) and 0.1 % formic acid in 80 % ACN (buffer B) at a flow rate of 400 nl/min with an effective gradient from 4 to 28 % of solvent B over 60 min, followed by 28 % to 44 % of buffer B over 13 min on an Vanquish Neo system (Thermo). Mass spectra were collected on an Orbitrap Eclipse mass spectrometer (Thermo). Full MS scans were performed in the mass analyzer over m/z range of 350-1,200 at a mass resolution of 60,000, while the MS/MS scans were implemented in the orbitrap mass analyzer with an isolation window of 1.6 m/z by the quadrupole mass filter at a mas resolution of 15,000. The data-dependent acquisition method was used in the cycle time mode with a cycle time of 2 s. The normalized collision energy of high collision density fragmentation was set to 30 and the dynamic exclusion time was set to 40 s.

For K27-IgG denaturing IP, peptides were separated on a self-packed C18 reversed-phase analytical column (75 µm × 20 cm, 1.9 µm) at a flow rate of 300 nL/min. The 60-minute liquid chromatography gradient was set as follows: 2-6 % phase B for 2 min, 6-22 % phase B from 2-49 min, 22-35 % phase B from 49-56.5 min, and 35-90 % phase B from 56.5-57 min, followed by maintenance at 90 % phase B until 60 min (phase A: 0.1 % formic acid in water; phase B: 99.9 % acetonitrile with 0.1 % formic acid). The mass spectrometer was operated in DIA-PASEF mode with an ion mobility range (1/K_0_) of 0.6-1.6 and a m/z scan range of 400-1201, using 32 scan windows and a ramp time of 100 ms. Ion mobility was calibrated using the Agilent ESI-L Low Concentration Tuning Mix.

For identification of the linkage type in UBE2Q1-assembled diUb, sample analysis was performed using a nano-liquid chromatography system (Easy nLC1200) coupled to a hybrid ultra-high-resolution mass spectrometer (Q Exactive plus, Thermo Fisher Scientific). Peptide samples were first separated same as K27-IgG denaturing IP. The mass spectrometer was operated in positive ion ESI mode using a data-dependent acquisition (DDA) method. Full MS scans were acquired in the Orbitrap mass analyzer at a resolution of 70,000 over a scan range of m/z 350-1,800, with an AGC target of 3e6 and a maximum injection time of 50 ms. For MS/MS analysis, precursor ions with charge states of 2-6 were selected and fragmented by higher-energy collisional dissociation (HCD) with a normalized collision energy (NCE) of 28.0 %. The secondary mass spectra were acquired at a resolution of 17,500 with an AGC target of 1e5 and a maximum injection time of 45 ms; dynamic exclusion was set to 30 s.

### Mass spectrometry data analysis and quantification

For K27-IgG native immunoprecipitation, raw data from LC-MS/MS measurements were analyzed using MaxQuant v2.4.2.0^79^ with default settings and match between runs and label-free quantification (LFQ) enabled. For protein identification, the human reference proteome downloaded from the UniProt (download date 2023/09/20, 20594 counts). FDR was determined by a reversed database and was set to 0.01 for peptides and proteins. Perseus software^80^ (version 1.6.7.0) was used for downstream statistical analysis. The volcano plots were created in the Perseus software and evaluated by two-sided Student’s t test with a permutation-based false discovery rate (FDR) control of 5 % and *s_0_* value of 0.1. Proteins identified with at least two unique peptides and has at least three valid quantification values in at least one group were taken into consideration for further data analysis. The missing values were replaced by random numbers draw from normal distribution (width = 0.3, downshift = 1.8). Log2 LFQ intensities were z-scored for heatmap visualization and hierarchical clustering. The protein-protein network was analyzed with STRING^81^. The gene ontology analysis was performed with DAVID bioinformatics^82^.

For K27-IgG denaturing IP, the DIA raw data were analyzed with Spectronaut 20 (Biognosis) using its factory default settings. The FDR threshold for protein qualification was also set to 1 %. Protein abundance was quantitatively integrated from the identified peptides using the MaxLFQ algorithm, and data were corrected using a local normalization strategy.

For identification of the linkage type in UBE2Q1-assembled diUb, the raw data were processed using PEAKS Studio 12.5 (Bioinformatics Solutions Inc.) and searched against the UniProt database. Mass tolerances for precursor and fragment ions were set to 10 ppm and 0.02 Da, respectively. The enzyme was set to trypsin, with a maximum of two missed cleavages allowed. Peptide and protein identification were filtered to a 1 % FDR. Label-free relative quantification was performed based on the peak intensity of the Top3 unique peptides, with normalization performed using the total ion current (TIC).

### Circular dichroism spectroscopy

Proteins were diluted with ultrapure water to a final concentration of 0.1 mg/mL. Circular dichroism spectra were acquired using a J-1500 CD Spectrometer (JASCO) at room temperature. The spectra of the proteins were measured three times from 195 nm to 260 nm in a 2 mm quartz cuvette at room temperature. Buffer baseline was subtracted from all spectra prior to analysis.

### CRISPR/Cas9-mediated DNAJB1 knockout

The construct PX459-sgDNAJB1 was transfected into HEK-293T cells, followed by selection with 1 μg/mL puromycin (Beyotime, ST551) for at least 6 days post-transfection. Surviving cells were subjected to limiting dilution in 96-well plates to isolate single-cell clones. Knockout was confirmed by western blot.

### Luciferase-based chaperone activity assay

Luciferase inactivation assay was performed according to previously protocol^83^. Recombinant firefly luciferase (50 nM; Sigma, SRE0045) was incubated at 42 °C alone or in the presence of the indicated proteins in luciferase refolding buffer (25 mM HEPES-KOH, pH 7.4, 150 mM potassium acetate,10 mM magnesium acetate and 10 mM DTT) supplemented with ATP-regeneration system (5 mM ATP, 30 mM creatine phosphate (Sigma, 27920) and 50 μg/mL creatine phosphokinase (Sigma, C3755), 10 mM magnesium chloride). The concentration of HSPA1B, DNAJB1 and K27-tetraUb was kept constant at 1 μM. Luciferase activities were measured using the Luciferase Assay System (Promega, E1500) in a microplate reader (BMG, CLARIOstar Plus) by mixing 5 μl of sample to 45 μl of luciferase assay reagent in white 96-well plates (Beyotime, FCP968). Luciferase refolding assay was performed following previously protocol^84^. Briefly, recombinant firefly luciferase (0.2 μM) was denatured at 30 °C for 40 min in denaturing buffer (6 M GuHCl, 25 mM HEPES-KOH, pH 7.5, 50 mM KCl, 10 mM MgCl_2_, 2 mM DTT). Refolding was initiated by 150-fold dilution into refolding buffer (25 mM HEPES-KOH, pH 7.5, 50 mM KCl, 10 mM MgCl_2_, 1 mM DTT, ATP-regeneration system) supplemented with the indicated proteins and incubated at 30 °C. Luminescence was measured at 0, 30, 60, 120 and 240 min as described above. The concentration of HSPA1B, DNAJB1 and K27-tetraUb was kept constant at 1 μM.

### ThT-based hIAPP fibrillization assay

Aggregation of hIAPP (10 µM) was carried out in aggregation assay buffer (50 mM HEPES, pH 7.4, 50 mM KCl, 2 mM TCEP) with 30 µM of indicated proteins. 10 µM of Thioflavin T (ThT; MCE, HY-D0218) was added to monitor aggregation kinetics. Reactions were assembled in 384-well plates (Thermo, 142761) and incubated at 37 °C in microplate reader without shaking. ThT fluorescence (Ex/Em = 430/485 nm) was measured every 10 minutes to monitor fibril formation. Experiments were performed in quadruplicate.

## Statistics

All experiments were independently repeated at least three times unless otherwise indicated. Statistical significance was evaluated using Student’s t-test or ANOVA, with *p*-value < 0.05 considered significant. No data were excluded and no statistical methods were used to predetermine sample size. Details are provided in the figure legends.

## References

1. Komander, D., and Rape, M. (2012). The Ubiquitin Code. Annu. Rev. Biochem. 81, 203–229. 10.1146/annurev-biochem-060310-170328.

2. Dikic, I., and Schulman, B.A. (2023). An expanded lexicon for the ubiquitin code. Nat. Rev. Mol. Cell Biol. 24, 273–287. 10.1038/s41580-022-00543-1.

3. Yau, R., and Rape, M. (2016). The increasing complexity of the ubiquitin code. Nat. Cell Biol. 18, 579–586. 10.1038/ncb3358.

4. Swatek, K.N., and Komander, D. (2016). Ubiquitin modifications. Cell Res. 26, 399–422. 10.1038/cr.2016.39.

5. De La Peña, A.H., Goodall, E.A., Gates, S.N., Lander, G.C., and Martin, A. (2018). Substrate-engaged 26 *S* proteasome structures reveal mechanisms for ATP-hydrolysis–driven translocation. Science 362, eaav0725. 10.1126/science.aav0725.

6. Rahman, S., and Wolberger, C. (2024). Breaking the K48-chain: linking ubiquitin beyond protein degradation. Nat. Struct. Mol. Biol. 31, 216–218. 10.1038/s41594-024-01221-w.

7. Marino, A., Di Fraia, D., Panfilova, D., Sahu, A.K., Minetti, A., Omrani, O., Cirri, E., and Ori, A. (2025). Aging and diet alter the protein ubiquitylation landscape in the mouse brain. Nat Commun 16, 5266. 10.1038/s41467-025-60542-6.

8. Ding, Z., Xu, C., Sahu, I., Wang, Y., Fu, Z., Huang, M., Wong, C.C.L., Glickman, M.H., and Cong, Y. (2019). Structural Snapshots of 26S Proteasome Reveal Tetraubiquitin-Induced Conformations. Mol. Cell 73, 1150–1161.e6. 10.1016/j.molcel.2019.01.018.

9. Dong, Y., Zhang, S., Wu, Z., Li, X., Wang, W.L., Zhu, Y., Stoilova-McPhie, S., Lu, Y., Finley, D., and Mao, Y. (2019). Cryo-EM structures and dynamics of substrate-engaged human 26S proteasome. Nature 565, 49–55. 10.1038/s41586-018-0736-4.

10. Cao, L., Liu, X., Zheng, B., Xing, C., and Liu, J. (2022). Role of K63-linked ubiquitination in cancer. Cell Death Discov 8, 410. 10.1038/s41420-022-01204-0.

11. Madiraju, C., Novack, J.P., Reed, J.C., and Matsuzawa, S. (2022). K63 ubiquitination in immune signaling. Trends Immunol. 43, 148–162. 10.1016/j.it.2021.12.005.

12. Takahashi, D.T., Wollscheid, H.-P., Lowther, J., and Ulrich, H.D. (2020). Effects of chain length and geometry on the activation of DNA damage bypass by polyubiquitylated PCNA. Nucleic Acids Res 48, 3042–3052. 10.1093/nar/gkaa053.

13. Wang, G., Gao, Y., Li, L., Jin, G., Cai, Z., Chao, J.-I., and Lin, H.-K. (2012). K63-Linked Ubiquitination in Kinase Activation and Cancer. Front. Oncol. 2. 10.3389/fonc.2012.00005.

14. Tracz, M., and Bialek, W. (2021). Beyond K48 and K63: non-canonical protein ubiquitination. Cell. Mol. Biol. Lett. 26, 1. 10.1186/s11658-020-00245-6.

15. Yu, Y., Zheng, Q., Erramilli, S.K., Pan, M., Park, S., Xie, Y., Li, J., Fei, J., Kossiakoff, A.A., Liu, L., et al. (2021). K29-linked ubiquitin signaling regulates proteotoxic stress response and cell cycle. Nat. Chem. Biol. 17, 896–905. 10.1038/s41589-021-00823-5.

16. Nucifora, F.C., Nucifora, L.G., Ng, C.-H., Arbez, N., Guo, Y., Roby, E., Shani, V., Engelender, S., Wei, D., Wang, X.-F., et al. (2016). Ubiqutination via K27 and K29 chains signals aggregation and neuronal protection of LRRK2 by WSB1. Nat. Commun. 7, 11792. 10.1038/ncomms11792.

17. Zhang, Q., Teng, X., Dai, Y., Wu, Y., Hou, H., and Li, J. (2025). K29-Linked Ubiquitination of Transcription Regulators Controls Cell Proliferation in the Unfolded Protein Response. Advanced Science, e09817. 10.1002/advs.202509817.

18. Michel, M.A., Elliott, P.R., Swatek, K.N., Simicek, M., Pruneda, J.N., Wagstaff, J.L., Freund, S.M.V., and Komander, D. (2015). Assembly and Specific Recognition of K29- and K33-Linked Polyubiquitin. Mol. Cell 58, 95–109. 10.1016/j.molcel.2015.01.042.

19. Garadi Suresh, H., Bonneil, E., Albert, B., Dominique, C., Costanzo, M., Pons, C., Masinas, M.P.D., Shuteriqi, E., Shore, D., Henras, A.K., et al. (2024). K29-linked free polyubiquitin chains affect ribosome biogenesis and direct ribosomal proteins to the intranuclear quality control compartment. Molecular Cell 84, 2337–2352.e9. 10.1016/j.molcel.2024.05.018.

20. Liu, J., Han, C., Xie, B., Wu, Y., Liu, S., Chen, K., Xia, M., Zhang, Y., Song, L., Li, Z., et al. (2014). Rhbdd3 controls autoimmunity by suppressing the production of IL-6 by dendritic cells via K27-linked ubiquitination of the regulator NEMO. Nat. Immunol. 15, 612–622. 10.1038/ni.2898.

21. Xue, B., Li, H., Guo, M., Wang, J., Xu, Y., Zou, X., Deng, R., Li, G., and Zhu, H. (2018). TRIM21 Promotes Innate Immune Response to RNA Viral Infection through Lys27-Linked Polyubiquitination of MAVS. J. Virol. 92, e00321–18. 10.1128/JVI.00321-18.

22. Lee, Y.R., Chen, M., Lee, J.D., Zhang, J., Lin, S.-Y., Fu, T.-M., Chen, H., Ishikawa, T., Chiang, S.-Y., Katon, J., et al. (2019). Reactivation of PTEN tumor suppressor for cancer treatment through inhibition of a MYC-WWP1 inhibitory pathway. Science 364, eaau0159. 10.1126/science.aau0159.

23. Liu, C., Liu, W., Ye, Y., and Li, W. (2017). Ufd2p synthesizes branched ubiquitin chains to promote the degradation of substrates modified with atypical chains. Nat. Commun. 8, 14274. 10.1038/ncomms14274.

24. Kulathu, Y., and Komander, D. (2012). Atypical ubiquitylation — the unexplored world of polyubiquitin beyond Lys48 and Lys63 linkages. Nat. Rev. Mol. Cell Biol. 13, 508–523. 10.1038/nrm3394.

25. Karim, M., Biquand, E., Declercq, M., Jacob, Y., Van Der Werf, S., and Demeret, C. (2020). Nonproteolytic K29-Linked Ubiquitination of the PB2 Replication Protein of Influenza A Viruses by Proviral Cullin 4-Based E3 Ligases. mBio 11, e00305–20. 10.1128/mBio.00305-20.

26. Kristariyanto, Y.A., Abdul Rehman, S.A., Campbell, D.G., Morrice, N.A., Johnson, C., Toth, R., and Kulathu, Y. (2015). K29-Selective Ubiquitin Binding Domain Reveals Structural Basis of Specificity and Heterotypic Nature of K29 Polyubiquitin. Mol. Cell 58, 83–94. 10.1016/j.molcel.2015.01.041.

27. Prus, G., Satpathy, S., Weinert, B.T., Narita, T., and Choudhary, C. (2024). Global, site-resolved analysis of ubiquitylation occupancy and turnover rate reveals systems properties. Cell 187, 2875–2892.e21. 10.1016/j.cell.2024.03.024.

28. Tsuchiya, H., Burana, D., Ohtake, F., Arai, N., Kaiho, A., Komada, M., Tanaka, K., and Saeki, Y. (2018). Ub-ProT reveals global length and composition of protein ubiquitylation in cells. Nat. Commun. 9, 524. 10.1038/s41467-018-02869-x.

29. Castañeda, C.A., Dixon, E.K., Walker, O., Chaturvedi, A., Nakasone, M.A., Curtis, J.E., Reed, M.R., Krueger, S., Cropp, T.A., and Fushman, D. (2016). Linkage via K27 Bestows Ubiquitin Chains with Unique Properties among Polyubiquitins. Structure 24, 423–436. 10.1016/j.str.2016.01.007.

30. Pan, M., Gao, S., Zheng, Y., Tan, X., Lan, H., Tan, X., Sun, D., Lu, L., Wang, T., Zheng, Q., et al. (2016). Quasi-Racemic X-ray Structures of K27-Linked Ubiquitin Chains Prepared by Total Chemical Synthesis. J. Am. Chem. Soc. 138, 7429–7435. 10.1021/jacs.6b04031.

31. Pei, G., Buijze, H., Liu, H., Moura-Alves, P., Goosmann, C., Brinkmann, V., Kawabe, H., Dorhoi, A., and Kaufmann, S.H.E. (2017). The E3 ubiquitin ligase NEDD4 enhances killing of membrane-perturbing intracellular bacteria by promoting autophagy. Autophagy 13, 2041–2055. 10.1080/15548627.2017.1376160.

32. Geisler, S., Holmström, K.M., Skujat, D., Fiesel, F.C., Rothfuss, O.C., Kahle, P.J., and Springer, W. (2010). PINK1/Parkin-mediated mitophagy is dependent on VDAC1 and p62/SQSTM1. Nat. Cell Biol. 12, 119–131. 10.1038/ncb2012.

33. Liu, Z., Chen, P., Gao, H., Gu, Y., Yang, J., Peng, H., Xu, X., Wang, H., Yang, M., Liu, X., et al. (2014). Ubiquitylation of Autophagy Receptor Optineurin by HACE1 Activates Selective Autophagy for Tumor Suppression. Cancer Cell 26, 106–120. 10.1016/j.ccr.2014.05.015.

34. Palicharla, V.R., and Maddika, S. (2015). HACE1 mediated K27 ubiquitin linkage leads to YB-1 protein secretion. Cell. Signal. 27, 2355–2362. 10.1016/j.cellsig.2015.09.001.

35. Ikeda, H., and Kerppola, T.K. (2008). Lysosomal localization of ubiquitinated jun requires multiple determinants in a lysine-27–linked polyubiquitin conjugate. Mol Biol Cell 19, 4588–4601. 10.1091/mbc.E08-05-0496.

36. Pan, X., Sun, Y., Liu, J., Chen, R., Zhang, Z., Li, C., Yao, H., and Ma, J. (2025). A bacterial RING ubiquitin ligase triggering stepwise degradation of BRISC via TOLLIP-mediated selective autophagy manipulates host inflammatory response. Autophagy 21, 1353–1372. 10.1080/15548627.2025.2468140.

37. Jin, S., Tian, S., Luo, M., Xie, W., Liu, T., Duan, T., Wu, Y., and Cui, J. (2017). Tetherin Suppresses Type I Interferon Signaling by Targeting MAVS for NDP52-Mediated Selective Autophagic Degradation in Human Cells. Mol. Cell 68, 308–322.e4. 10.1016/j.molcel.2017.09.005.

38. Cao, Z., Conway, K.L., Heath, R.J., Rush, J.S., Leshchiner, E.S., Ramirez-Ortiz, Z.G., Nedelsky, N.B., Huang, H., Ng, A., Gardet, A., et al. (2015). Ubiquitin Ligase TRIM62 Regulates CARD9-Mediated Anti-fungal Immunity and Intestinal Inflammation. Immunity 43, 715–726. 10.1016/j.immuni.2015.10.005.

39. Sparrer, K.M.J., Gableske, S., Zurenski, M.A., Parker, Z.M., Full, F., Baumgart, G.J., Kato, J., Pacheco-Rodriguez, G., Liang, C., Pornillos, O., et al. (2017). TRIM23 mediates virus-induced autophagy via activation of TBK1. Nature Microbiology 2, 1543–1557. 10.1038/s41564-017-0017-2.

40. Gatti, M., Pinato, S., Maiolica, A., Rocchio, F., Prato, M.G., Aebersold, R., and Penengo, L. (2015). RNF168 Promotes Noncanonical K27 Ubiquitination to Signal DNA Damage. Cell Rep. 10, 226–238. 10.1016/j.celrep.2014.12.021.

41. El-Hachem, N., Habel, N., Naiken, T., Bzioueche, H., Cheli, Y., Beranger, G.E., Jaune, E., Rouaud, F., Nottet, N., Reinier, F., et al. (2018). Uncovering and deciphering the pro-invasive role of HACE1 in melanoma cells. Cell Death & Differentiation 25, 2010–2022. 10.1038/s41418-018-0090-y.

42. Yin, Q., Han, T., Fang, B., Zhang, G., Zhang, C., Roberts, E.R., Izumi, V., Zheng, M., Jiang, S., Yin, X., et al. (2019). K27-linked ubiquitination of BRAF by ITCH engages cytokine response to maintain MEK-ERK signaling. Nat. Commun. 10, 1870. 10.1038/s41467-019-09844-0.

43. Cunningham, C.N., Baughman, J.M., Phu, L., Tea, J.S., Yu, C., Coons, M., Kirkpatrick, D.S., Bingol, B., and Corn, J.E. (2015). USP30 and parkin homeostatically regulate atypical ubiquitin chains on mitochondria. Nat. Cell Biol. 17, 160–169. 10.1038/ncb3097.

44. Swatek, K.N., Usher, J.L., Kueck, A.F., Gladkova, C., Mevissen, T.E.T., Pruneda, J.N., Skern, T., and Komander, D. (2019). Insights into ubiquitin chain architecture using Ub-clipping. Nature 572, 533–537. 10.1038/s41586-019-1482-y.

45. Singh Gautam, A.K., and Matouschek, A. (2019). Decoding without the cipher. Nat. Chem. Biol. 15, 210–212. 10.1038/s41589-019-0230-9.

46. Miura, G. (2017). Unraveling the chain. Nat. Chem. Biol. 13, 1139–1139. 10.1038/nchembio.2508.

47. Newton, K., Matsumoto, M.L., Wertz, I.E., Kirkpatrick, D.S., Lill, J.R., Tan, J., Dugger, D., Gordon, N., Sidhu, S.S., Fellouse, F.A., et al. (2008). Ubiquitin Chain Editing Revealed by Polyubiquitin Linkage-Specific Antibodies. Cell 134, 668–678. 10.1016/j.cell.2008.07.039.

48. Matsumoto, M.L., Wickliffe, K.E., Dong, K.C., Yu, C., Bosanac, I., Bustos, D., Phu, L., Kirkpatrick, D.S., Hymowitz, S.G., Rape, M., et al. (2010). K11-Linked Polyubiquitination in Cell Cycle Control Revealed by a K11 Linkage-Specific Antibody. Mol. Cell 39, 477–484. 10.1016/j.molcel.2010.07.001.

49. Matsumoto, M.L., Dong, K.C., Yu, C., Phu, L., Gao, X., Hannoush, R.N., Hymowitz, S.G., Kirkpatrick, D.S., Dixit, V.M., and Kelley, R.F. (2012). Engineering and Structural Characterization of a Linear Polyubiquitin-Specific Antibody. J. Mol. Biol. 418, 134–144. 10.1016/j.jmb.2011.12.053.

50. Michel, M.A., Swatek, K.N., Hospenthal, M.K., and Komander, D. (2017). Ubiquitin Linkage-Specific Affimers Reveal Insights into K6-Linked Ubiquitin Signaling. Mol. Cell 68, 233–246.e5. 10.1016/j.molcel.2017.08.020.

51. Yau, R.G., Doerner, K., Castellanos, E.R., Haakonsen, D.L., Werner, A., Wang, N., Yang, X.W., Martinez-Martin, N., Matsumoto, M.L., Dixit, V.M., et al. (2017). Assembly and Function of Heterotypic Ubiquitin Chains in Cell-Cycle and Protein Quality Control. Cell 171, 918–933.e20. 10.1016/j.cell.2017.09.040.

52. Lange, S.M., McFarland, M.R., Lamoliatte, F., Carroll, T., Krshnan, L., Pérez-Ràfols, A., Kwasna, D., Shen, L., Wallace, I., Cole, I., et al. (2024). VCP/p97-associated proteins are binders and debranching enzymes of K48–K63-branched ubiquitin chains. Nat Struct Mol Biol 31, 1872–1887. 10.1038/s41594-024-01354-y.

53. Pan, M., Zheng, Q., Ding, S., Zhang, L., Qu, Q., Wang, T., Hong, D., Ren, Y., Liang, L., Chen, C., et al. (2019). Chemical Protein Synthesis Enabled Mechanistic Studies on the Molecular Recognition of K27-linked Ubiquitin Chains. Angew Chem Int Ed 58, 2627–2631. 10.1002/anie.201810814.

54. Fang, G.-M., Li, Y.-M., Shen, F., Huang, Y.-C., Li, J.-B., Lin, Y., Cui, H.-K., and Liu, L. (2011). Protein chemical synthesis by ligation of peptide hydrazides. Angew. Chem. Int. Ed. 50, 7645–7649. 10.1002/anie.201100996.

55. Zheng, J.-S., Tang, S., Qi, Y.-K., Wang, Z.-P., and Liu, L. (2013). Chemical synthesis of proteins using peptide hydrazides as thioester surrogates. Nat. Protoc. 8, 2483–2495. 10.1038/nprot.2013.152.

56. Miller, K.R., Koide, A., Leung, B., Fitzsimmons, J., Yoder, B., Yuan, H., Jay, M., Sidhu, S.S., Koide, S., and Collins, E.J. (2012). T Cell Receptor-Like Recognition of Tumor In Vivo by Synthetic Antibody Fragment. PLoS ONE 7, e43746. 10.1371/journal.pone.0043746.

57. Paduch, M., Koide, A., Uysal, S., Rizk, S.S., Koide, S., and Kossiakoff, A.A. (2013). Generating conformation-specific synthetic antibodies to trap proteins in selected functional states. Methods 60, 3–14. 10.1016/j.ymeth.2012.12.010.

58. Chen, Z., and Pickart, C.M. (1990). A 25-kilodalton ubiquitin carrier protein (E2) catalyzes multi-ubiquitin chain synthesis via lysine 48 of ubiquitin. J. Biol. Chem. 265, 21835–21842.

59. Hofmann, R.M., and Pickart, C.M. (1999). Noncanonical MMS2-Encoded Ubiquitin-Conjugating Enzyme Functions in Assembly of Novel Polyubiquitin Chains for DNA Repair. Cell 96, 645–653. 10.1016/s0092-8674(00)80575-9.

60. Qu, Q., Pan, M., Gao, S., Zheng, Q., Yu, Y., Su, J., Li, X., and Hu, H. (2018). A Highly Efficient Synthesis of Polyubiquitin Chains. Adv. Sci. 5, 1800234. 10.1002/advs.201800234.

61. Tang, S., Liang, L., Si, Y., Gao, S., Wang, J., Liang, J., Mei, Z., Zheng, J., and Liu, L. (2017). Practical Chemical Synthesis of Atypical Ubiquitin Chains by Using an Isopeptide-Linked Ub Isomer. Angew Chem Int Ed 56, 13333–13337. 10.1002/anie.201708067.

62. Maine, G.N., Li, H., Zaidi, I.W., Basrur, V., Elenitoba-Johnson, K.S.J., and Burstein, E. (2010). A bimolecular affinity purification method under denaturing conditions for rapid isolation of a ubiquitinated protein for mass spectrometry analysis. Nat Protoc 5, 1447–1459. 10.1038/nprot.2010.109.

63. Cheng, X., Wang, Y., Liu, J., Wu, Y., Zhang, Z., Liu, H., Tian, L., Zhang, L., Chang, L., Xu, P., et al. (2024). Super Enhanced Purification of Denatured-Refolded Ubiquitinated Proteins by ThUBD Revealed Ubiquitinome Dysfunction in Liver Fibrosis. Molecular & Cellular Proteomics 23, 100852. 10.1016/j.mcpro.2024.100852.

64. McClellan, A.J., Tam, S., Kaganovich, D., and Frydman, J. (2005). Protein quality control: chaperones culling corrupt conformations. Nat Cell Biol 7, 736–741. 10.1038/ncb0805-736.

65. Wickner, S., Maurizi, M.R., and Gottesman, S. (1999). Posttranslational Quality Control: Folding, Refolding, and Degrading Proteins. Science 286, 1888–1893. 10.1126/science.286.5446.1888.

66. Rosenzweig, R., Nillegoda, N.B., Mayer, M.P., and Bukau, B. (2019). The Hsp70 chaperone network. Nat. Rev. Mol. Cell Biol. 20, 665–680. 10.1038/s41580-019-0133-3.

67. Ayala Mariscal, S.M., Pigazzini, M.L., Richter, Y., Özel, M., Grothaus, I.L., Protze, J., Ziege, K., Kulke, M., ElBediwi, M., Vermaas, J.V., et al. (2022). Identification of a HTT-specific binding motif in DNAJB1 essential for suppression and disaggregation of HTT. Nat Commun 13, 4692. 10.1038/s41467-022-32370-5.

68. Hageman, J., Rujano, M.A., van Waarde, M.A.W.H., Kakkar, V., Dirks, R.P., Govorukhina, N., Oosterveld-Hut, H.M.J., Lubsen, N.H., and Kampinga, H.H. (2010). A DNAJB chaperone subfamily with HDAC-dependent activities suppresses toxic protein aggregation. Mol. Cell 37, 355–369. 10.1016/j.molcel.2010.01.001.

69. Monistrol, J., Beton, J.G., Johnston, E.C., Dang, T.L., Bukau, B., and Saibil, H.R. (2025). Stepwise recruitment of chaperone Hsc70 by DNAJB1 produces ordered arrays primed for bursts of amyloid fibril disassembly. Commun Biol 8, 522. 10.1038/s42003-025-07906-2.

70. Wentink, A.S., Nillegoda, N.B., Feufel, J., Ubartaitė, G., Schneider, C.P., De Los Rios, P., Hennig, J., Barducci, A., and Bukau, B. (2020). Molecular dissection of amyloid disaggregation by human HSP70. Nature 587, 483–488. 10.1038/s41586-020-2904-6.

71. Qin, F., Cao, R., Cui, W., Bai, X., Yuan, J., Zhang, Y., Liu, Y., Cao, N., Dong, N., Zhou, M., et al. (2025). Listerin promotes α-synuclein degradation to alleviate Parkinson’s disease through the ESCRT pathway. Sci. Adv. 11, eadp3672. 10.1126/sciadv.adp3672.

72. Rho, H., Kim, S., Kim, S.U., Kim, J.W., Lee, S.H., Park, S.H., Escorcia, F.E., Chung, J.-Y., and Song, J. (2024). CHIP ameliorates nonalcoholic fatty liver disease via promoting K63- and K27-linked STX17 ubiquitination to facilitate autophagosome-lysosome fusion. Nat. Commun. 15, 8519. 10.1038/s41467-024-53002-0.

73. Lord, D.M., Bird, J.J., Honey, D.M., Best, A., Park, A., Wei, R.R., and Qiu, H. (2018). Structure-based engineering to restore high affinity binding of an isoform-selective anti-TGFβ1 antibody. mAbs 10, 444–452. 10.1080/19420862.2018.1426421.

74. Michel, M.A., Komander, D., and Elliott, P.R. (2018). Enzymatic Assembly of Ubiquitin Chains. In The Ubiquitin Proteasome System: Methods and Protocols, T. Mayor and G. Kleiger, eds. (Springer), pp. 73–84. 10.1007/978-1-4939-8706-1_6.

75. Kabsch, W. (2010). XDS. Acta Crystallogr D Biol Crystallogr 66, 125–132. 10.1107/S0907444909047337.

76. Emsley, P., and Cowtan, K. (2004). *Coot*: model-building tools for molecular graphics. Acta Crystallogr D Biol Crystallogr 60, 2126–2132. 10.1107/S0907444904019158.

77. Agirre, J., Atanasova, M., Bagdonas, H., Ballard, C.B., Baslé, A., Beilsten-Edmands, J., Borges, R.J., Brown, D.G., Burgos-Mármol, J.J., Berrisford, J.M., et al. (2023). The *CCP* 4 suite: integrative software for macromolecular crystallography. Acta Crystallogr D Struct Biol 79, 449–461. 10.1107/S2059798323003595.

78. Pettersen, E.F., Goddard, T.D., Huang, C.C., Meng, E.C., Couch, G.S., Croll, T.I., Morris, J.H., and Ferrin, T.E. (2021). UCSF CHIMERAX: Structure visualization for researchers, educators, and developers. Protein Science 30, 70–82. 10.1002/pro.3943.

79. Cox, J., and Mann, M. (2008). MaxQuant enables high peptide identification rates, individualized p.p.b.-range mass accuracies and proteome-wide protein quantification. Nat Biotechnol 26, 1367–1372. 10.1038/nbt.1511.

80. Tyanova, S., Temu, T., Sinitcyn, P., Carlson, A., Hein, M.Y., Geiger, T., Mann, M., and Cox, J. (2016). The Perseus computational platform for comprehensive analysis of (prote)omics data. Nat Methods 13, 731–740. 10.1038/nmeth.3901.

81. Szklarczyk, D., Kirsch, R., Koutrouli, M., Nastou, K., Mehryary, F., Hachilif, R., Gable, A.L., Fang, T., Doncheva, N.T., Pyysalo, S., et al. (2023). The STRING database in 2023: protein–protein association networks and functional enrichment analyses for any sequenced genome of interest. Nucleic Acids Res 51, D638–D646. 10.1093/nar/gkac1000.

82. Sherman, B.T., Hao, M., Qiu, J., Jiao, X., Baseler, M.W., Lane, H.C., Imamichi, T., and Chang, W. (2022). DAVID: a web server for functional enrichment analysis and functional annotation of gene lists (2021 update). Nucleic Acids Res 50, W216–W221. 10.1093/nar/gkac194.

83. Huang, L., Agrawal, T., Zhu, G., Yu, S., Tao, L., Lin, J., Marmorstein, R., Shorter, J., and Yang, X. (2021). DAXX represents a new type of protein-folding enabler. Nature 597, 132–137. 10.1038/s41586-021-03824-5.

84. Faust, O., Abayev-Avraham, M., Wentink, A.S., Maurer, M., Nillegoda, N.B., London, N., Bukau, B., and Rosenzweig, R. (2020). HSP40 proteins use class-specific regulation to drive HSP70 functional diversity. Nature 587, 489–494. 10.1038/s41586-020-2906-4.

85. Ma, J., Chen, T., Wu, S., Yang, C., Bai, M., Shu, K., Li, K., Zhang, G., Jin, Z., He, F., et al. (2019). iProX: An integrated proteome resource. Nucleic Acids Res. 47, D1211–D1217. 10.1093/nar/gky869.

86. Chen, T., Ma, J., Liu, Y., Chen, Z., Xiao, N., Lu, Y., Fu, Y., Yang, C., Li, M., Wu, S., et al. (2022). iProX in 2021: Connecting proteomics data sharing with big data. Nucleic Acids Res. 50, D1522–D1527. 10.1093/nar/gkab1081.

