## Supplemental information for "Conformation-specific Antibody Deciphers K27-linked Ubiquitination in Chaperone-Mediated Proteostasis"

**Figure S1-S9**

**Table S1**

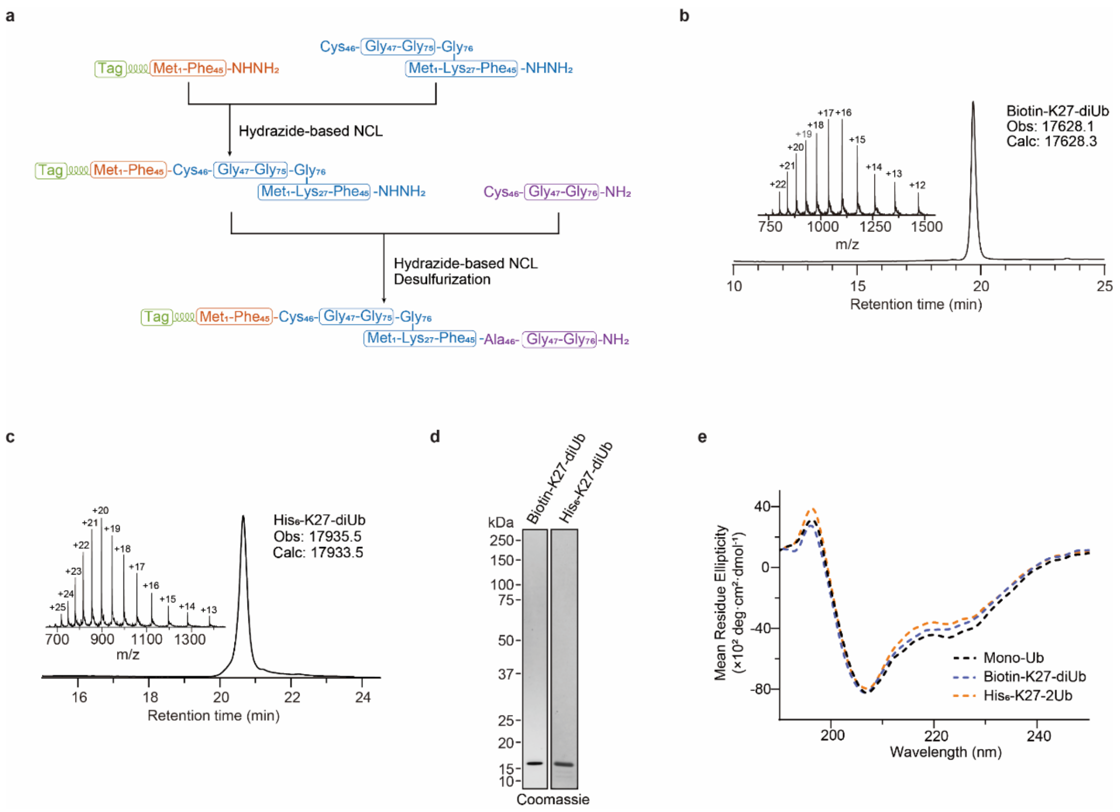

**Figure S1. Characterization of synthetic K27-linked diUb.**

**(a)** Schematic of the synthetic strategy used to generate tagged K27-linked diUb. **(b-c)** Analytical LC-MS confirming the identity and purity of synthetic K27-linked diUb. **(d)** SDS-PAGE analysis of synthetic K27-linked diUbs. **(e)** CD spectra comparing refolded K27-linked diUb and monoUb.

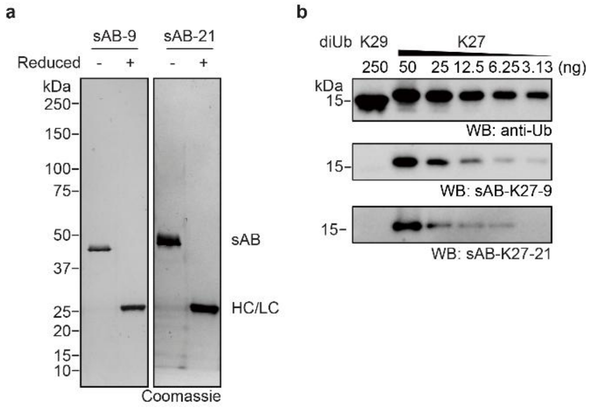

**Figure S2. Characterization and comparison of K27 linkage-specific antibodies.**
**(a)** SDS–PAGE analysis of purified sABs. HC, heavy chain. LC, light chain. **(b)** Western blot showing specificity of sAB-K27 for K27-linked diUb against K29-linked diUb. All gel panels in this figure are representative at least three independent experiments.

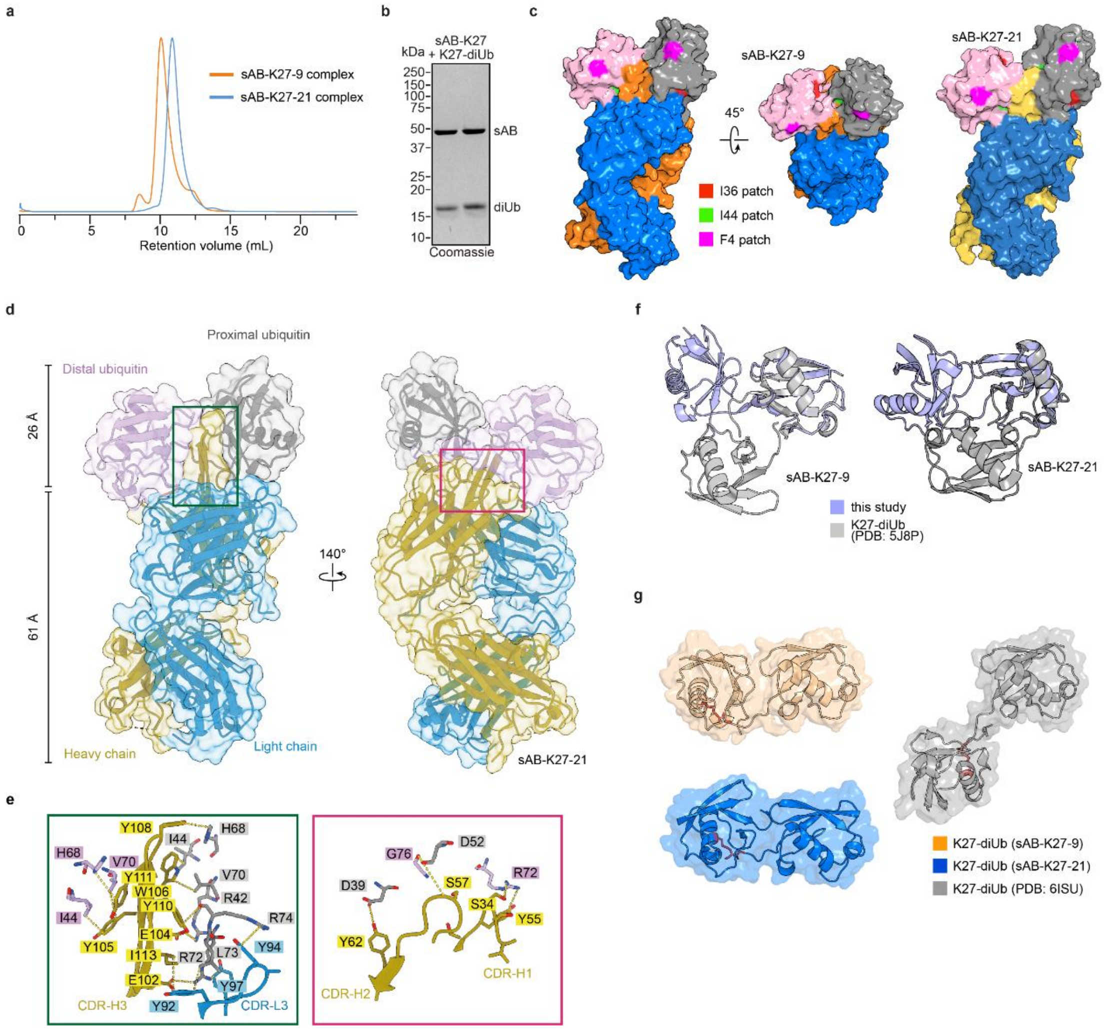

**Figure S3. Purification, crystallization, and structural comparisons of K27-linked diUb in complex with sAB-K27.**

**(a)** Size-exclusion chromatography of purified sAB-K27 in complex with K27-linked diUb. **(b)** SDS–PAGE analysis confirming purity of the purified complexes and demonstrating a homogeneous 1:1 stoichiometry. **(c)** Structural analysis showing sAB-K27 recognition of the hydrophobic patches of K27-linked diUb, notably the Ile44 patch. **(d)** Same as in Fig. 2a but with sAB-K27-21, resolved at 2.9 Å. **(e)** Same as in Fig. 2b but with sAB-K27-21. **(f)** Structural comparison between the K27-linked diUb in this study and the apo form (PDB 5J8P). Distal ubiquitin molecules were aligned. **(g)** Structural comparison between the K27-linked diUb in this study and the K27-linked diUb in complex with UCHL3 (PDB 6ISU). Distal ubiquitin molecules were aligned.

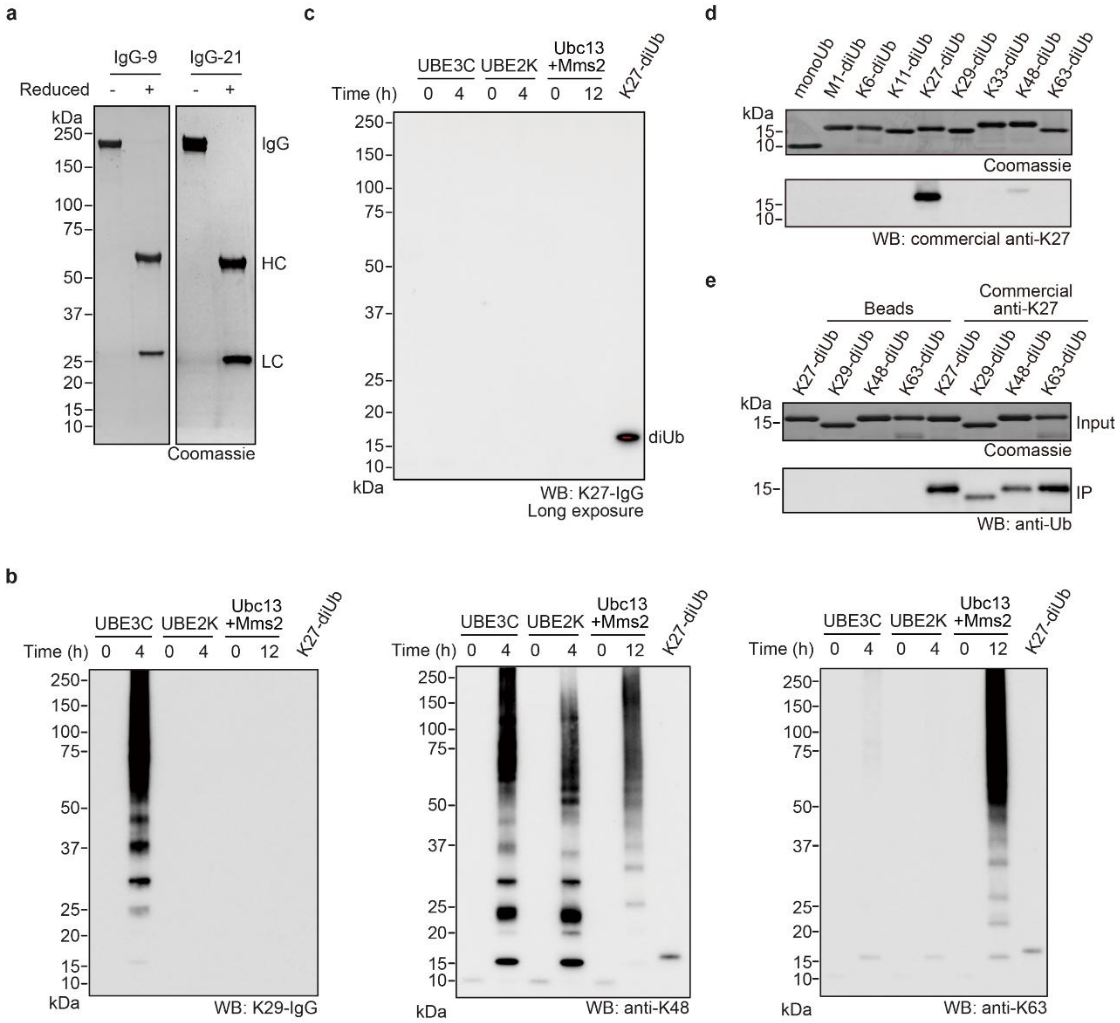

**Figure S4. Characterization and comparison of K27 linkage-specific antibodies.** **(a)** SDS-PAGE analysis of purified K27-IgGs. HC, heavy chain. LC, light chain. **(b)** Western blot of *in vitro* assembled polyubiquitin chains using ubiquitin linkage-specific (K29, K48 and K63) antibodies. **(c)** Western blot demonstrating the specificity of K27-IgG for K27-linked diUb over K29-, K48-, and K63-linked polyubiquitin. Overexposure is indicated in red. **(d-e)** Evaluation of the commercial sequence-specific K27 antibody by western blotting (d) and immunoprecipitation (e), revealing limited specificity and narrow applicability. Gel panels in (b) and (c) are representative at least two independent experiments.

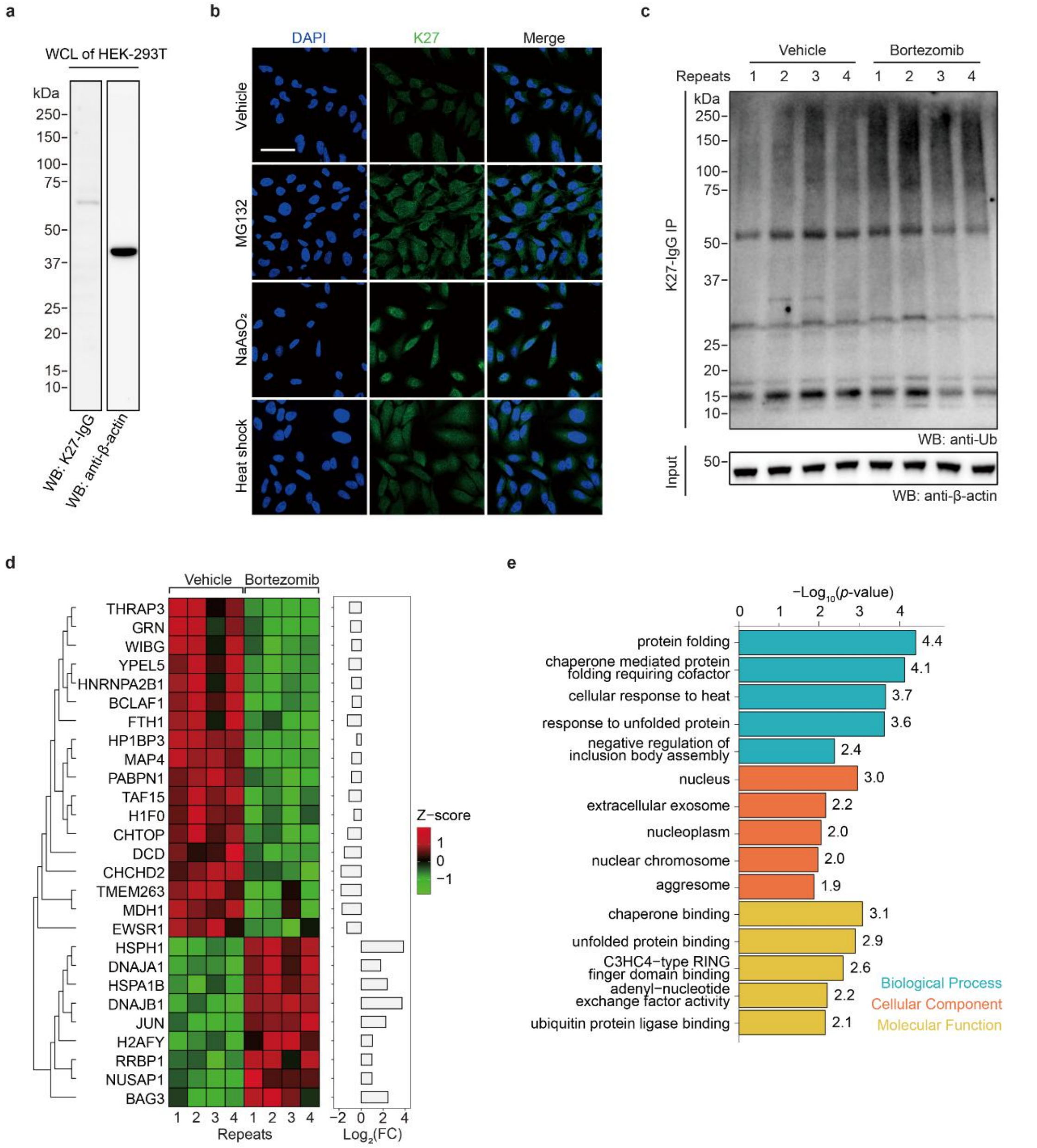

**Figure S5. Proteomic profiling of bortezomib-treated cells by native IP using K27-IgG.**

**(a)** Direct Western blot analysis of K27-linked ubiquitination in HEK-293T cells treated with bortezomib (1 µM, 4 h), MG132 (10 µM, 4 h), sodium arsenite (250 µM, 4 h), or heat shock (43 °C, 30 min) without immunoprecipitation enrichment. **(b)** Immunofluorescence imaging of K27-linked ubiquitin (green) in HeLa cells treated with MG132 (10 µM, 4 h), sodium arsenite (250 µM, 4 h), or heat shock (43 °C, 30 min). Scale bar, 50 µm. **(c)** Western blot validation of K27-IgG native IP from HEK-293T cells treated with bortezomib (1 µM, 4 h). **(d)** Heat map showing upregulated and downregulated proteins identified by proteomic profiling under bortezomib treatment. **(e)** GO analysis of upregulated proteins, revealing strong enrichment with protein folding and molecular chaperone activity. All gel panels and images in this figure are representative at least two independent experiments.

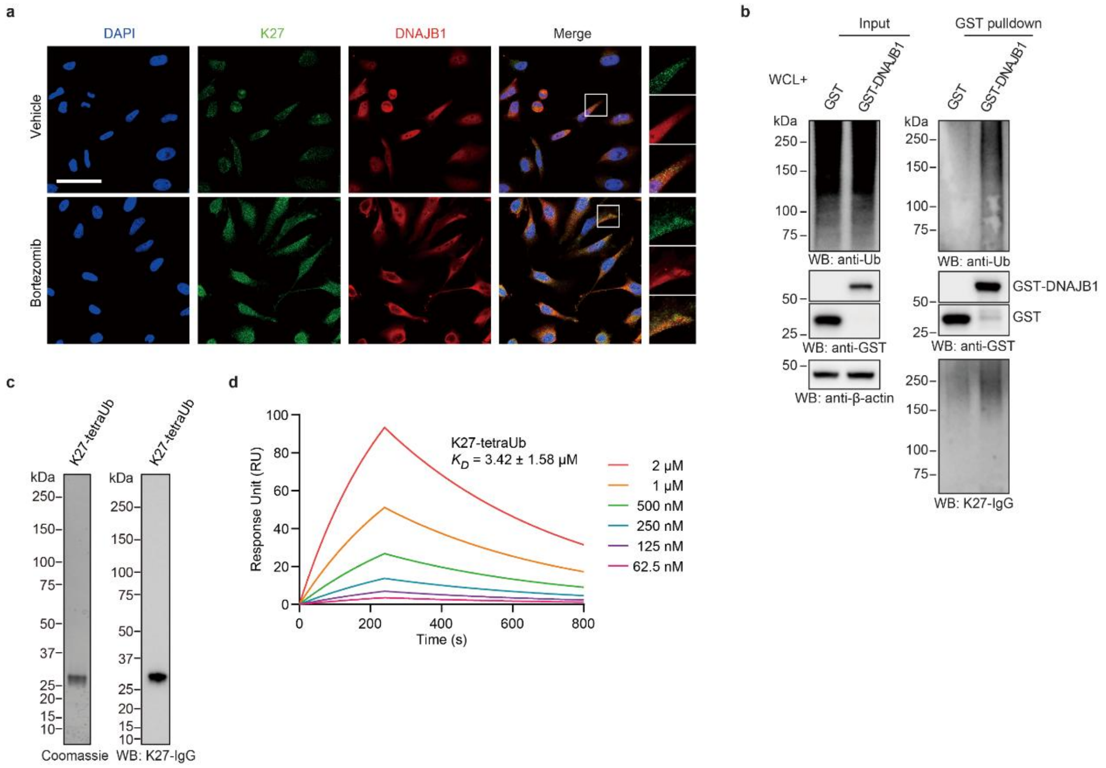

**Figure S6. K27-linked ubiquitination interacts with DNAJB1.**

**(a)** Immunofluorescent imaging of HeLa cells treated with bortezomib (1 μM, 4 h), showing partial co-localization of DNAJB1 (red) and K27-linked ubiquitin (green). Scale bar, 50 μm. **(b)** Western blot of K27-linked polyubiquitin co-immunoprecipitated with purified GST-DNAJB1 from HEK293T cells treated with bortezomib (1 μM, 4 h) using K27-IgG. GST alone was used as a control. **(c)** Western blot analysis of K27-tetraUb using K27-IgG. **(d)** SPR analysis showing DNAJB1 binds to synthetic K27-linked tetraUb with measurable $K_D$ of 3.42 ± 1.58 μM. All gel panels and images in this figure are representative at least two independent experiments.

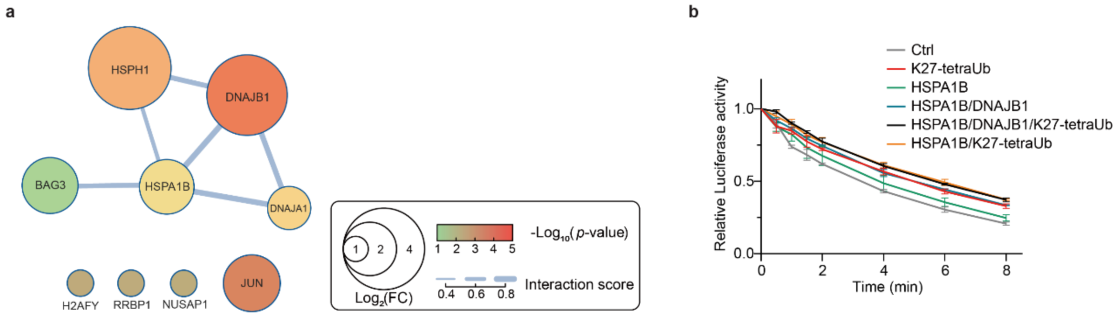

**Figure S7. K27-linked ubiquitin modulates the HSP70 chaperone system and potentially suppresses protein aggregation.**

**(a)** Protein-protein interaction network of the significantly enriched proteins upon bortezomib treatment. **(b)** Kinetic curves of luciferase inactivation assays showing that K27-linked tetraUb (1 μM) synergizes with the HSP70-DNAJB1 complex to prevent thermal inactivation. K27-linked ubiquitin alone also delays inactivation, indicating a chaperone-independent protective effect. Data are presented relative to the initial value (n = 3).

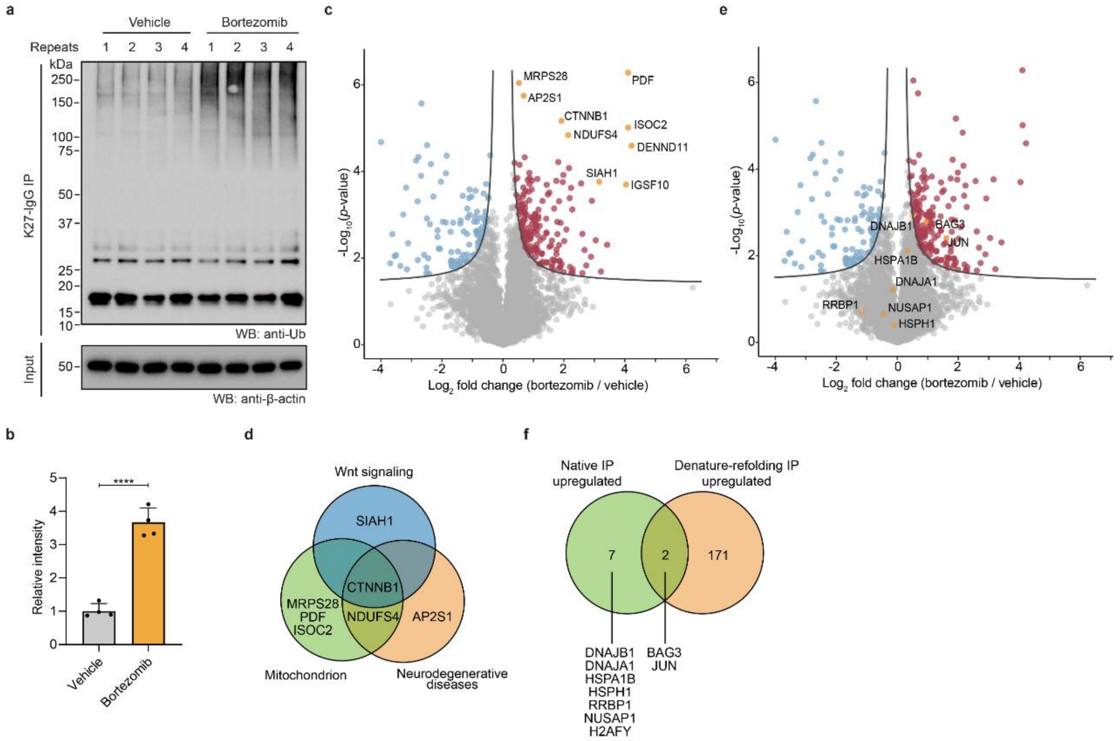

**Figure S8. Proteomic profiling of bortezomib-treated cells by DRUSP-based IP using K27-IgG.**

**(a-b)** Bortezomib-treated (1 µM, 4 h) HEK293T cell lysates were subjected to DRUSP-based denaturing immunoprecipitation with K27-IgG. (n = 4, biological replicates; one-tailed Student's t-test; ****, $p < 0.0001$) **(c)** Volcano plot of differentially enriched proteins identified by proteomic analysis (n = 4, biological replicates). Significantly enriched proteins (FDR < 0.05, s0 = 0.1) are shown in red, and depleted proteins are in blue, the top enriched proteins are indicated in orange. **(d)** Venn plot classifying the top enriched proteins by their known functions in mitochondrion, neurodegenerative diseases and Wnt signaling. **(e)** Volcano plot as in (c), the proteins enriched in native IP are indicated in orange. **(f)** Venn plot comparing the proteins enriched by native IP versus DRUSP-based IP.

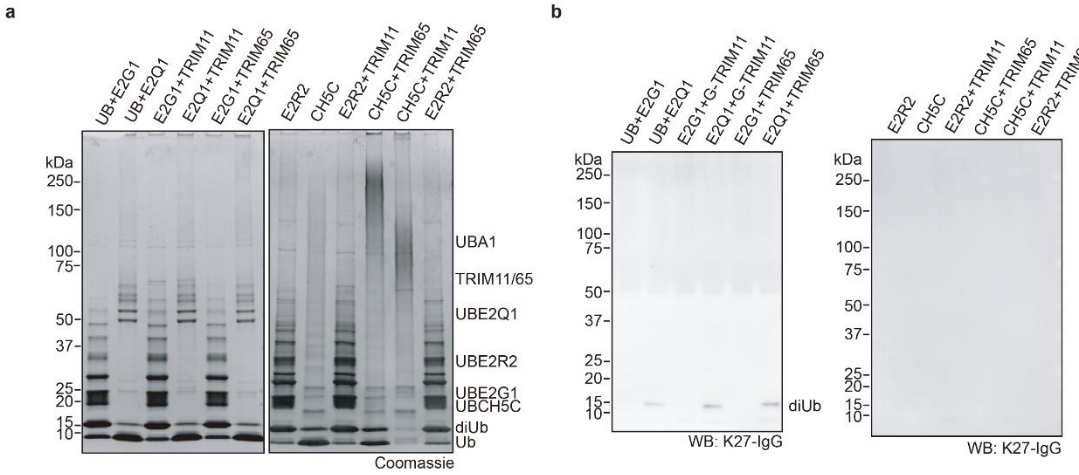

**Figure S9. *In vitro* ubiquitination assays using E2 and E3 enzymes identified by denaturing IP.**

**(a)** *In vitro* ubiquitination assays using E2 and E3 enzymes identified by denaturing IP, including UBE2G1, UBE2R2 and TRIM11, TRIM65. UBE2G1 and UBCH5C were tested since they are known E2 enzymes for TRIM E3 ligase family. **(b)** Western blot analysis of reaction products from (a) using K27-IgG revealing that UBE2Q1 is potentially capable of forming K27-linked diUb. All gel panels and images in this figure are representative at least two independent experiments.

**Table S1. X-ray data collection and refinement statistics, related to Figure 2**

| | sAB-K27-9&K27-diUb complex | sAB-K27-21&K27-diUb complex |
|---|---|---|
| **Data collection** | | |
| Space group | $P2_1$ | $P\ 2_1\ 2_1\ 2_1$ |
| Cell dimensions | | |
| *a*, *b*, *c* (Å) | 92.34, 71.50, 93.91 | 78.74, 137.91, 145.79 |
| α, β, γ (°) | 90, 95.08, 90 | 90, 90, 90 |
| Resolution (Å) | 2.2 (2.26-2.20) * | 2.9 (2.9-3.0) |
| $R_{sym}$ or $R_{merge}$ | 0.075 (0.85) | 0.06207 (0.8329) |
| $CC_{1/2}$ (%) | 99.8 (68.5) | 0.999 (0.664) |
| $I$ / σ$I$ | 9.75 (1.75) | 14.26 (1.44) |
| Completeness (%) | 99.2 (98.9) | 98.23 (98.50) |
| Redundancy | 3.8 (3.9) | 3.4(3.5) |
| **Refinement** | | |
| Resolution (Å) | 68.7 – 2.2 | 100.2 – 2.9 |
| No. reflections | 61663 | 35301 |
| $R_{work}$ / $R_{free}$ | 19.4 / 23.1 | 21.6/26.0 |
| No. atoms | | |
| Protein | 9115 | 9160 |
| Ligand/ion | 73 | 40 |
| Water | 340 | 37 |
| *B*-factors | | |
| Protein | 51.2 | 81.8 |
| Ligand/ion | 72.0 | 91.0 |
| Water | 50.4 | 69.6 |
| R.m.s. deviations | | |
| Bond lengths (Å) | 0.004 | 0.002 |
| Bond angles (°) | 0.654 | 0.53 |

*Number of xtals for each structure should be noted in footnote. *Values in parentheses are for highest-resolution shell.

[AU: Equations defining various *R*-values are standard and hence are no longer defined in the footnotes.]

[AU: Ramachandran statistics should be in Methods section at the end of Refinement subsection.]

[AU: Wavelength of data collection, temperature and beamline should all be in Methods section.]
